# Deficiency of the hemoglobin-haptoglobin receptor, CD163, worsens insulin sensitivity in obese male mice

**DOI:** 10.1101/2024.05.31.596887

**Authors:** Michael W. Schleh, Magdalene Ameka, Alec Rodriguez, Alyssa H. Hasty

## Abstract

Excessive iron accumulation in metabolic organs such as the adipose tissue, liver, and skeletal muscle is associated with increased diabetes risk. Tissue-resident macrophages serve multiple roles including managing inflammatory tone and regulating parachymal iron homeostasis; thus protecting against metabolic dysfunction upon iron overload. The scavenger receptor CD163 is uniquely present on tissue-resident macrophages, and plays a significant role in iron homeostasis by clearing extracellular hemoglobin-haptoglobin complexes, thereby limiting oxidative damage caused by free hemoglobin in metabolic tissues. We show that the absence of CD163 exacerbates glucose intolerance and insulin resistance in male mice with obesity. Additionally, loss of CD163 reduced the expression of iron regulatory genes *(Tfr1, Cisd1, Slc40a1)* in adipose tissue macrophages and anti-inflammatory (M2-like) bone marrow-derived macrophages (BMDMs). Further, CD163 deficiency mediated a pro-inflammatory shift and limited hemoglobin scavenging specifically in M2-like BMDMs. To this end, iron buffering was diminished in inguinal white adipose tissue (iWAT) macrophages *in vivo*, which culminated in iron spillover into adipocytes and CD45^+^CD11B^-^ non-myeloid immune cells in iWAT. These findings show that CD163 on tissue-resident macrophages is critical for their anti-inflammatory and hemoglobin scavenging roles, and its absence results in impaired systemic insulin action in an obese setting.

**Article Highlights:**

- Loss of CD163 mediates a phenotypic switch in M2-like macrophages towards a pro-inflammatory state.
- CD163 is involved in free hemoglobin uptake and catabolism as well as oxidative metabolism, specifically in M2-like macrophages.
- In inguinal white adipose tissue (iWAT) of CD163 defficient mice, macrophage iron is reduced; while concomitantly, adipocyte and other immune cell iron content is increased.
- Loss of CD163 provokes glucose intolerance and insulin resistance in obese male mice.

## Introduction

Maintaining proper iron balance is critical for metabolic health. Imbalanced tissue iron levels are linked to metabolic diseases such as nonalcoholic fatty liver disease and type 2 diabetes (1). Iron is essential for many cellular functions including ATP generation, DNA synthesis and repair, and oxygen transport. Therefore, various cellular and tissue-specific regulatory pathways are critical for tissue level and systemic iron homeostasis (2). The uptake of dietary iron is regulated by duodenal enterocytes, which engage with transferrin (Tf) for cellular uptake through the Tf receptor (Tfr) to support erythropoiesis (3). Iron is retained systemically through the phagocytosis of senescent erythrocytes by splenic red pulp macrophages leading to recycling of Tf-bound iron into the circulation (2). Additionally, tissue-resident macrophages are specialized with resolving inflammatory mediators in tissues caused by factors such as obesity and infection (4); and also facilitate storage or transfer of iron, thus contributing to the maintenance of local tissue iron homeostasis (5, 6).

Macrophages serve homeostasic needs of cells in response to environmental cues, and are often binned into two classifications: pro-inflammatory (M1-like) and anti-inflammatory (M2-like). However, this generic classification is governed by *in vitro* polarization states, and macrophage populations present *in vivo* span a variety of phenotypes and can simultaneously express M1 and M2 markers (7, 8). Macrophage phenotypes also underlie their iron handling capacity, where M2-like tissue-resident macrophages are noted for their intrinsic capacity to provide bioavailable iron within the tissue microenvironment (5, 9). Macrophages endocytose ferritin-bound iron via receptors such as Tfr proteins (TFR1/2), and iron-containing molecules such as hemoglobin-haptoglobin (Hb-Hp) and heme-hemopexin via the scavenger receptors CD163 and CD91, respectively. Importantly, CD163 serves as a cell-surface marker for tissue-resident macrophages (10). Therefore, the anti-inflammatory function of CD163 is defined by hemoglobin-bound iron uptake and catabolism, thereby limiting metabolic tissues from iron overload. Hence, the ability for tissue-resident macrophages to effectively respond to environmental stimuli while also fulfilling their anti-inflammatory functions is crucial for tissue homeostasis.

Metabolic complications in obesity are often associated with inflammatory activation of macrophages in metabolically-significant tissues such as adipose tissue (11, 12). Macrophages are the most abundant immune cell type in adipose tissue and can expand from ∼10% to ∼40% of all immune cells in the lean versus obese states (13, 14). The majority of macrophages recruited to obese adipose tissue are pro-inflammatory, and release cytokines such as IL-1β, TNFα, and IL-6 (13-15). These cytokines are noted to promote adipose tissue insulin resistance and increase lipolysis (16). During inflammation, macrophage iron handling capacity is also forfeited (9, 17). In obesity, the presence of iron-handling CD163^+^ tissue-resident macrophages is diminished in favor of macrophages specialized in managing the lipid rich environment [e.g., recruitment of lipid-associated macrophages; (18)]. This shift in macrophage phenotypes may explain why adipocytes are susceptible to iron overload in obesity (17, 19). Adipocyte iron overload is speculated to initiate the temporal response to insulin resistance by inducing systemic dyslipidemia and reducing adiponectin release (19-22). Therefore, interventions aimed to counter the relative loss of Cd163^+^ tissue-resident macrophages may be therapeutically advantageous to limit excessive free hemoglobin, iron overload, and inflammatory stress caused by obesity.

Here, we report loss of CD163 results in adipocyte iron overload in inguinal white adipose tissue (iWAT) in high-fat diet (HFD) fed male mice. Furthermore, CD163 deficiency hindered the polarization of macrophages towards an M2-like phenotype and reduced hemoglobin scavenging *in vitro*. Overall, these studies suggest that CD163^+^ tissue-resident macrophages are critical in preventing severe metabolic dysfunction caused by diet-induced obesity, preserving an anti-inflammatory iron handling phenotype in M2-like macrophages, and limiting adipocyte iron overload.

## Research Design and Methods

### Mouse Model

Experimental protocols were approved by the Vanderbilt University Institutional Animal Care and Use Committee. Male and female Cd163^-/-^ mice on C57BL/6N background were purchased from Jackson Laboratory (Cd163^tm1.1(KOMP)Vlcg^) and backcrossed to 99% purity with the C57BL/6J background. CD163^+/-^ mice were bred, and CD163^-/-^ or CD163^+/+^ (WT littermates) were randomized to either low-fat diet (LFD; Research Diets, 10% fat, 3.85 kcal/g) or HFD (Research Diets, 60% fat, 5.24 kcal/g) for 16 weeks. All mice were fed *ad libitum* in a 12 h light/dark cycle at 22°C.

### Body composition

Fat-free mass (FFM) and fat mass (FM) were measured from conscious mice at 16 weeks diet by nuclear magnetic resonance (Bruker Minispec).

### Intraperitoneal Glucose Tolerance Testing

The day after body composition assessments, intraperitoneal glucose tolerance tests (ipGTTs) were performed (2 g/kg FFM, 20% dextrose). Mice were fasted for 4 h, and blood glucose was collected using a Contour Next EZ blood glucose monitoring system at 0, 15, 30, 45, 60, 90, 120 min following glucose injection. Glucose area under the curve was calculated using the trapezoidal rule.

### Hyperinsulinemic-euglycemic clamp

Hyperinsulinemic-euglycemic clamp studies were performed by the Vanderbilt Mouse Metabolic Phenotyping Center. Prodecures are detailed in **Supplementary Methods** and at https://vmmpc.org.

### Adipocyte and stromal cell isolation

The isolation of epididymal white adipose tissue (eWAT) and iWAT adipocytes from the stromal vascular fraction (SVF) was completed by digesting in 2 mg/mL collagenase type IV (Worthington), as detailed in the **Supplementary Methods** and previously (23, 24).

### Flow cytometry and fluorescence-activated cell sorting

Adipose tissue SVF cells were analyzed and sorted on a FACS Aria III cell sorter (BD Biosciences), at the Vanderbilt University Medical Center Flow Cytometry Shared Resource Core. A complete list of antibody is provided in **Supplementary Table 1.** Cells from eWAT and iWAT were blocked for 30 min at 4°C in anti-mouse CD16/CD32 Fc Block, then stained for 30 min at 4°C with following antibodies; BV510 anti-CD45, PerCP/Cyanine5.5 anti-CD11b, APC/Cyanine7 F4/80. DAPI was used for viability. Stained cells were then washed twice with PBS + 1% FBS and isolated by fluorescence-activated cell sorting (FACS) into 15 mL metal-free conical tubes. Samples were suspended in 200 μL 70% HNOL for inductively coupled plasma mass spectrometry (ICP-MS) analysis.

### Bone marrow derived macrophages

Male and female mice fed chow diets were euthanized between 8-16 weeks of age. Bone-marrow derived macrophages (BMDMs) were extracted from the femurs and tibia, using a 22-gauge needle to flush marrow using cold RPMI. After suspension, cells were treated with red blood cell lysis buffer, and neutralized with PBS + 1% FBS. Cells were plated in BMDM media for 7 days (∼5×10^6^ cells per flask). Following 7 days differentiation, M1-like BMDMs were treated with 10 ng/mL LPS + 100 ng/mL IFNγ for 24 h and M2-like BMDMs were treated with 10 ng/mL IL-4 + 10 ng/mL IL-13 for 72 h.

### RNA isolation, reverse transcription and quantitative RT–PCR

Tissues or cells were lysed, DNase treated, and RNA was purified using a Qiagen RNeasy Mini Kit (Qiagen), then reverse transcribed into cDNA (iScript RT; BioRad). RT-qPCR was performed using FAM-conjugated TaqMan Gene Expression Assay primers (ThermoFisher) and iQ Supermix (BioRad). Samples were normalized to GAPDH or β-actin and quantified by the 2^-ΔΔCT^ method. Complete lists of primers are in **Supplementary Table 2**.

### RNA Sequencing

RNA from unpolarized (M0), and M1- or M2-like polarized BMDMs were purified with the RNeasy Plus Mini Kit (Qiagen). Library preparation included Poly-A enrichment using Poly- A magnetic beads (New England Biolabs, E776), and cDNA library prep using the NEBNext Ultra Kit (New England Biolabs, E7760). Paired-end sequencing was completed using the Illumina NovaSeq 6000 sequencing platform targeting an average of 50 M reads per sample.

Differential expression for individual gene read counts were analyzed with DESeq2 1.36 (25), using R 4.3.1, and explained previously by our lab (26). Differential expression analyses between WT vs. Cd163^-/-^ BMDMs were corrected for multiple comparisons using Benjamini Hochberg methods (27).

### Histology

Tissue iron was visualized at 10X (Leica DMi8) using the Perls Prussian Blue perfusion and staining method described previously (17, 28). Following cardiac perfusion, ∼15 mL Prussian blue stain (4% paraformaldehyde, 1% potassium ferrocyanide, 1% HCl) was perfused and fixed for 1 h. Tissues were then incubated in Perls staining solution for 12 h at 4°C before paraffin embedding.

### ICP-MS

Tissues, adipocytes, and SVF populations were homogenized and digested by 70% HNO_3_ in metal-free conical tubes (VWR). Total iron was quantified by ICP-MS (Agilent), and detailed in **Supplementary Methods**.

### Mitochondrial respiratory capacity

M0, M1-, and M2-like BMDMs were plated in 96-well Seahorse assay plates (50,000-75,000/well) in Seahorse assay media, and mitochondrial respiration was analyzed on a Seahorse XFe96 Extracellular Flux Analyzer (Agilent). For mitochondria stress tests, BMDMs were sequentially treated with 10 mM glucose, 1 mM oligomycin, 1 mM FCCP, and 0.5 mM antimycin A + rotenone. Oxygen consumption rate (OCR) was normalized to cellular protein content.

### Statistics

Student’s *t*-tests were performed for between group measurements. For studies that included main effects of treatment or time (e.g., ipGTT and clamps), a 2-way ANOVA was completed, with Tukey multiple comparisons correction. RNA sequencing and figure generation was completed using R 4.3.1, and the remainder of analyses were performed by GraphPad Prism 10.1.2. Data are expressed as mean ± SEM, and significance set to α = 0.05.

## Results

### Loss of Cd163 impairs glucose tolerance in obesity

CD163 is expressed exclusively on human monocytes/macrophages (**Figure 1A**), and tissue-resident macrophages in mice (23). To examine the role of CD163 on metabolic homeostasis, a 16-week LFD and HFD feeding paradigm was implemented (**Figure 1B**). Deletion of CD163 was confirmed in both eWAT and iWAT adipose tissue macropahges (ATMs; **Figure 1C**). Body mass was tightly linked to diet (HFD vs. LFD), and body weight was similar between CD163^-/-^ mice and WT littermates (**Figure 1D**). Following 16 weeks on diet, male CD163^-/-^ mice on LFD had slightly lower FFM (p=0.04) and greater FM (p=0.02); while CD163^-/-^ mice on HFD had lower FFM compared to WT littermates (p=0.04; **Figure 1E**). No difference in daily energy intake was observed between CD163^-/-^ mice and WT littermates in either diet group (**Figure 1F**). In female mice, body weight did not differ between genotypes (**Supplementary Figure 1A**).

**Figure 1.**
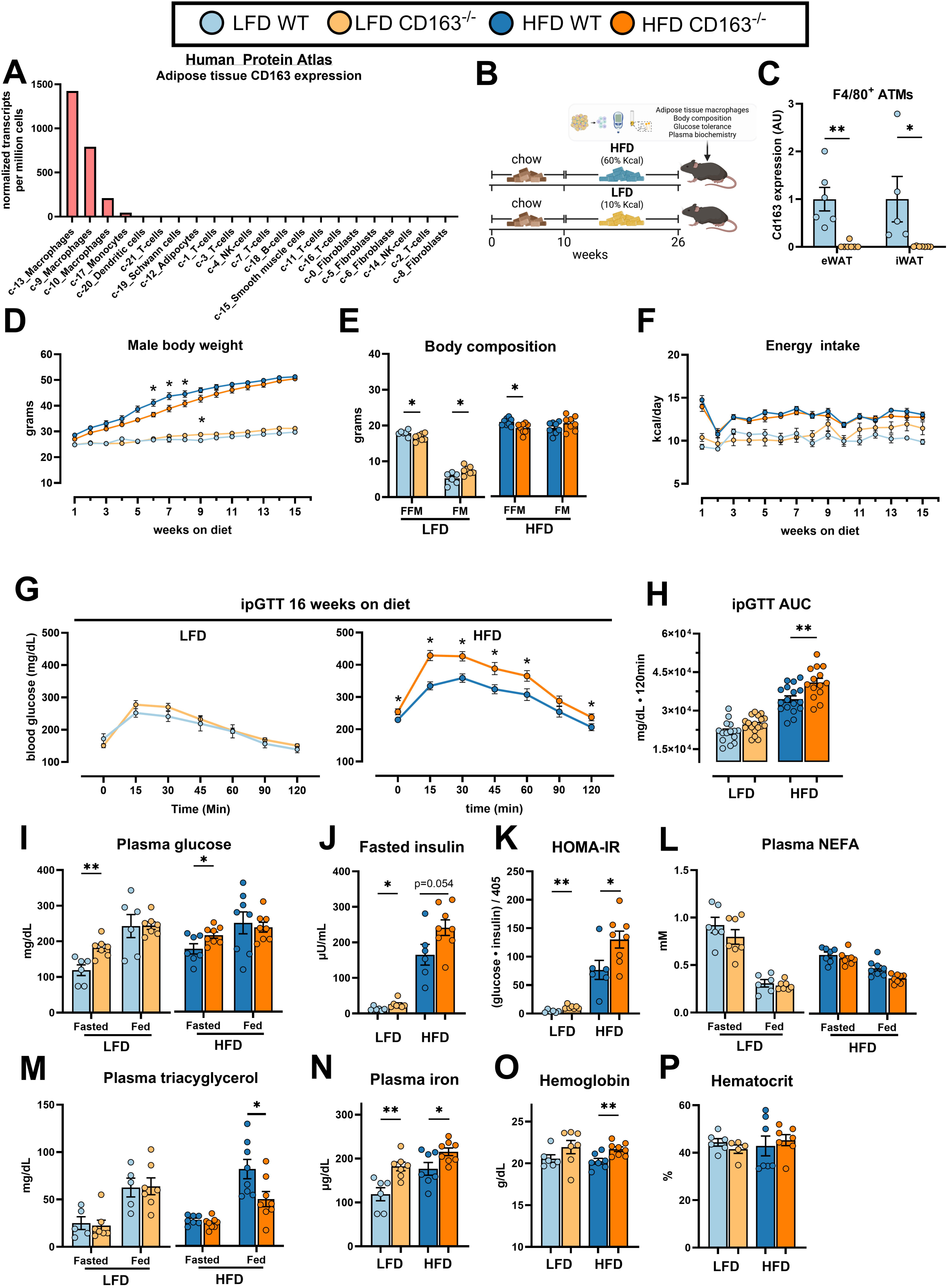
Loss of Cd163 impairs glucose tolerance in obesity. A) Human Protein Atlas single-cell transcriptomic analysis of adipose tissue CD163 expression (47). B) Study design; Male CD163^-/-^ mice and WT littermates were randomly selected to LFD or HFD groups for 16 weeks. C) ATM (F4/80^+^) expression of CD163 in eWAT and iWAT. D-F) Weekly body mass throughout the 16-week diet study (D), body composition at 16 weeks diet (E), and daily energy intake (kcal/day; F). G-H) ipGTT (G) and ipGTT AUC (H) performed after a 4 h fast at 16 weeks on either LFD or HFD. I-K) Plasma glucose measured in the fasted and fed state (I), fasted insulin (J), and HOMA-IR ([glucose (mg/dL) • insulin (µU/mL)] / 405; K). L-M) Plasma NEFA (L) and triacylglycerol (M) measured in the fasted and fed state. N-P) Plasma iron (N), whole blood hemoglobin (O), and hematocrit (P). Unpaired Student’s *t*-tests assessed differences between CD163^-/-^ vs. WT littermates on either LFD or HFD in panels C,E,H,I-P. Two-way repeated measures ANOVA assessed differences between genotypes over time for measures conducted in panels D, and F-G. Multiple comparisons were assessed with Tukey’s post hoc assessment. Data are presented as mean ± SEM of 8-17 mice per group. ATM, adipose tissue macrophage; AUC, area under the curve; eWAT, epididymal white adipose tissue; FFM, fat-free mass; FM, fat mass; HFD, high-fat diet; HOMA-IR, homeostatic model assessment for insulin resistance; ipGTT, intraperitoneal glucose tolerance test; iWAT, inguinal white adipose tissue; LFD, low-fat diet. *p<0.05, **p<0.01; genotype affect, WT vs. Cd163^-/-^.

While there was no difference in glucose tolerance by genotype in LFD or HFD males at 4, 8, or 12 weeks (**Supplementary Figure 2**); glucose tolerance was worsened in CD163^-/-^ males compared to WT controls after 16 weeks on HFD (**Figure 1G & H**). Female CD163^-/-^ and WT controls had similar glucose tolerance at all timepoints (**Supplementary Figure 1B-E**). At 16 weeks, blood glucose was similar between the genotypes in fed animals; however, fasting plasma glucose was higher in CD163^-/-^ males in lean and obese mice (**Figure 1I**). Fasting insulin was significantly greater in CD163^-/-^ males on LFD, and trended greater in the CD163^-/-^ on HFD (**Figure 1J**). The homeostatic model assessment for insulin resistance (HOMA-IR) is commonly used to assess insulin sensitivity in clinical settings (29), and was greater in the CD163^-/-^ mice on LFD and HFD (**Figure 1K**). Plasma non-esterified FA (NEFA) and triacylglycerol did not differ between genotypes in the fasted or fed state, apart from fed state triacylglycerol, which was lower in the CD163^-/-^ mice (**Figure 1L & M**). Due to the hemoglobin scavenging properties of CD163, the CD163^-/-^ mice on both LFD and HFD presented greater plasma iron (**Figure 1N**), while total hemoglobin in circulation was only greater in CD163^-/-^ mice on HFD (**Figures 1O**). Hematocrit did not differ between groups (**Figure 1P**). These data collectively show that loss of CD163 attenuates glucose tolerance and elevates circulating iron and hemoglobin in obesity.

### Impaired glucose tolerance by loss of CD163 is attributed to peripheral insulin resistance

To assess whether the impaired glucose tolerance in the CD163^-/-^ mice was attributed to peripheral insulin resistance, hyperinsulinemic-euglycemic clamps were conducted (**Supplementary Figure 3**). Data males are shown in **Figure 2**, and for females are shown in **Supplementary Figure 4**. Body mass did not differ between CD163^-/-^ male mice and WT littermates after 1-week catheterization (**Figure 2A**). As designed, insulin levels increased from the basal to clamp states (endogenous glucose production + exogenous insulin infusion), with no significant difference by genotype (**Figure 2B**). Arterial glucose levels were clamped at 150 mg/dL by regulating the variable glucose infusion rate [GIR; **Figure 2C**), and in HFD-fed mice, the GIR required to maintain euglycemia was nearly 2-fold lower in CD163^-/-^ mice compared to WT littermates (**Figure 2D**). Consistent with lower GIR necessary to maintain euglycemia, glucose rate of disappearance (Rd), calculated by the sum of GIR and glucose rate of appearance (Ra), was attenuated in CD163^-/-^ mice only on HFD (**Figure 2E**). Glucose Ra did not differ across groups in the basal or clamp states (**Figure 2F**). Similarly, glucose Ra suppression (index of hepatic insulin sensitivity) did not significantly differ across groups, although it trended lower in CD163^-/-^ mice (**Figure 2G**). CD163^-/-^ mice presented greater NEFA throughout the clamp on LFD and HFD (**Figure 2H & 2I**) suggesting resistance to the suppressive effects of insulin. NEFA percent suppression from basal to clamp conditions did not significantly differ by genotype in either diet condition (**Figure 2J**).

**Figure 2.**
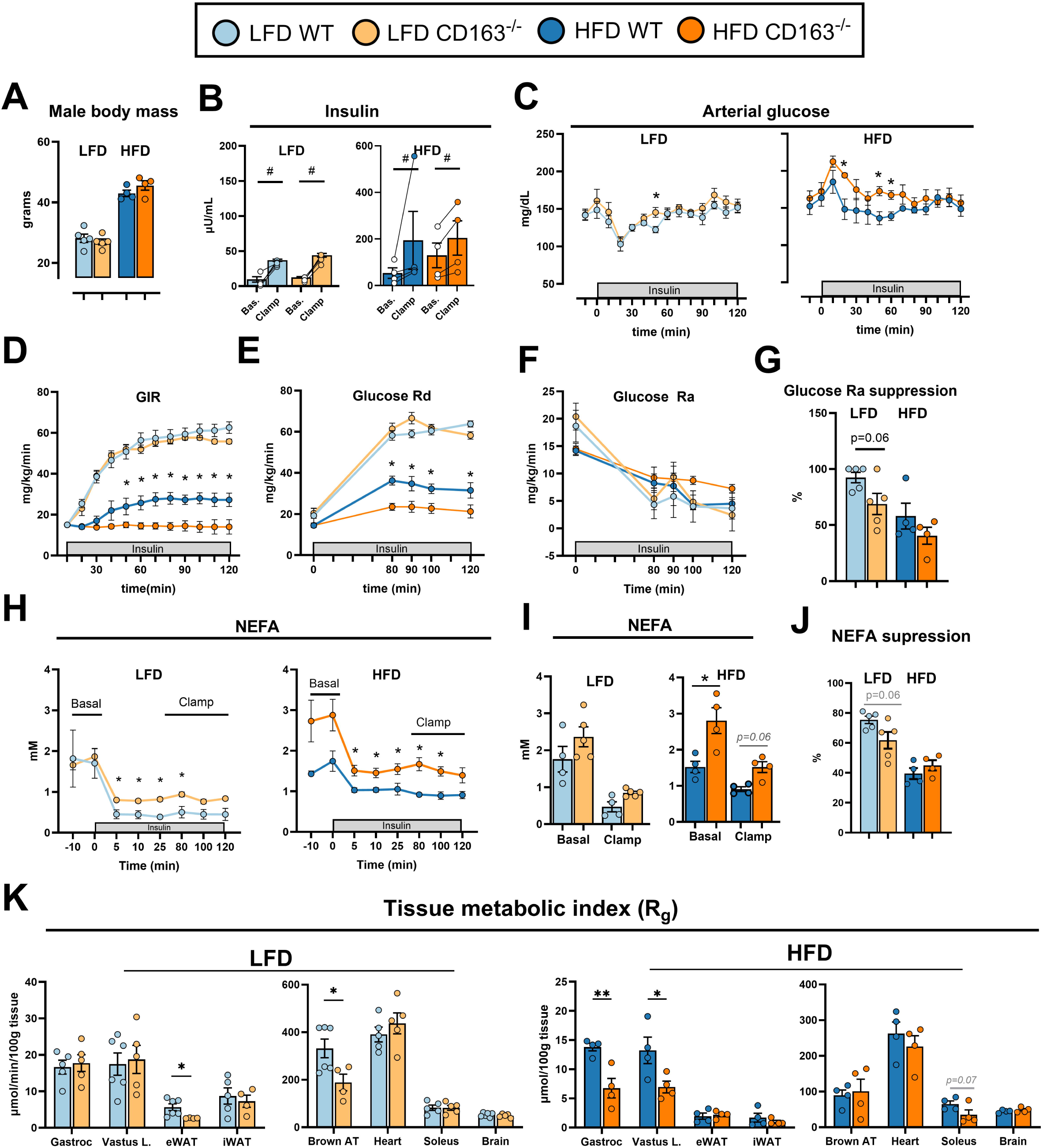
Impaired glucose tolerance by loss of CD163 is attributed to peripheral insulin resistance. A) Body weight was similar between CD163^-/-^ male mice and WT littermates following one-week catheter indwelling in both HFD and LFD groups. B) Insulin concentration increased from basal (t=-10 min) and clamped (t=80-120 min) conditions in all sampled mice. C) Arterial glucose was measured continuously with target concentration of 150 mg/dL. D) Exogenous GIR was controlled to maintain euglycemia in LFD and HFD mice. E) Glucose Rd, measured by the sum of exogenous GIR and endogenous glucose Ra. F) Glucose Ra measured during the hyperinsulinemic clamp. G) Glucose Ra percent suppression from basal to clamp conditions (index of hepatic insulin sensitivity). H-J) NEFA measured from arterial plasma during the clamp (H), during basal and clamp conditions (I), and NEFA percent suppression from basal to clamp conditions (J). K) A bolus of [^14^C]2DG was infused at 120 min. Mice were then anaesthetized and tissues snap frozen at 155 min to measure for ^14^C radioactivity (normalized per 100 g tissue) in LFD and HFD male mice. Unpaired Student’s *t*-test was used to compare differences in CD163^-/-^ and WT littermates in LFD and HFD conditions in panels A, G, J, and K. Two-way repeated measures ANOVA was used to assess differences between genotypes over time during the clamp, and conducted in figure panels B-F, and H-I. Pairwise comparisons were adjusted with Tukey correction. Data are expressed mean ± SEM of 4-6 mice per group. 2DG, 2-Deoxy-D-glucose; eWAT, epididymal white adipose tissue; Gastroc, gastrocnemius; HFD, high-fat diet; iWAT, inguinal white adipose tissue; LFD, low-fat diet; NEFA, non-esterified fatty acid; Ra, endogenous rate of glucose appearance; Rd, glucose rate of disappearance; Vastus L, vastus lateralis. *p<0.05, **p<0.01; genotype affect, WT vs. Cd163^-/-^. #p<0.05 main effect for time during the hyperinsulinemic clamp, basal vs. clamp.

Differences in glucose disposal are largely attributed to impaired glucose uptake from peripheral tissues. The tissue metabolic index (Rg) estimates the rate of glucose uptake in selected tissues, and the CD163^-/-^ mice on LFD presented lower eWAT and brown adipose tissue (BAT) Rg (**Figure 2K**). The CD163^-/-^ mice on HFD significantly decreased Rg in the gastrocnemius, vastus lateralis, and trended lower in the soleus (**Figure 2K**). Female CD163^-/-^ mice showed greater insulin concentrations during the hyperinsulinemic clamp only on HFD (**Supplementary Figure 4B**), but displayed no differences in GIR or glucose Rd, NEFA response to insulin, or R_g_ on either diet (**Supplementary Figure 4D-J**). These findings indicate that CD163^-/-^ male mice with obesity exhibit insulin resistance, elevations in circulating NEFA in the presence of high insulin, and decreased Rg in skeletal muscle.

### CD163 protects iWAT from iron overload in obesity

Total iron content (^56^Fe) was measured by ICP-MS in eWAT, iWAT, BAT, liver, gastrocnemius, heart, pancreas, thymus, and spleen. CD163^-/-^ mice on HFD had significantly greater iWAT and BAT tissue iron (**Figure 3A**). Further, adipocytes were isolated from the eWAT and iWAT (**Figure 3B**). Iron content eWAT adipocytes was not affected by CD163 deficiency or diet, while adipocytes from iWAT of CD163^-/-^ mice on HFD presented greater iron concentrations (**Figure 3C**). To assess the effects of CD163 on SVF immune cell iron handling, FACS was implemented to sort nonhematopoietic cells (CD45^-^), non-myeloid lymphocytes (CD45^+^CD11B^-^), and macrophages (CD45^+^CD11B^+^F4/80^+^; **Figure 3D**). Iron content in CD45^-^ cells was greater in iWAT versus eWAT, and significantly greater iron was detected in iWAT CD45^+^CD11B^-^ in CD163^-/-^ mice (**Figure 3E**). In contrast, iron in CD45^+^CD11B^+^F4/80^+^ ATMs from iWAT of CD163^-/-^ mice was lower than controls (**Figure 3E**). Additionally, ATMs were isolated from eWAT and iWAT by magnetic sorting (F4/80^+^ fraction), and gene expression for iron handling genes were analyzed by qPCR. Gene expression for *Tfr1* and *Cisd1* was lower in CD163^-/-^ mice in both eWAT and iWAT ATMs, whereas ferroportin (*Slc40a1*) and *Ncoa4* were decreased only in iWAT ATMs (**Figure 3F**). Prussian Blue staining did not display major visual differences in iron depositions or structure from the eWAT or iWAT, whereas BAT from CD163^-/-^ mice appeared more unilocular compared to WT littermates (**Figure 3G**). Altogether, these findings illustrate iron handling is altered in CD163^-/-^ iWAT, as described by decreased ATM iron content and a conseuqnetial increase in adipocyte iron.

**Figure 3.**
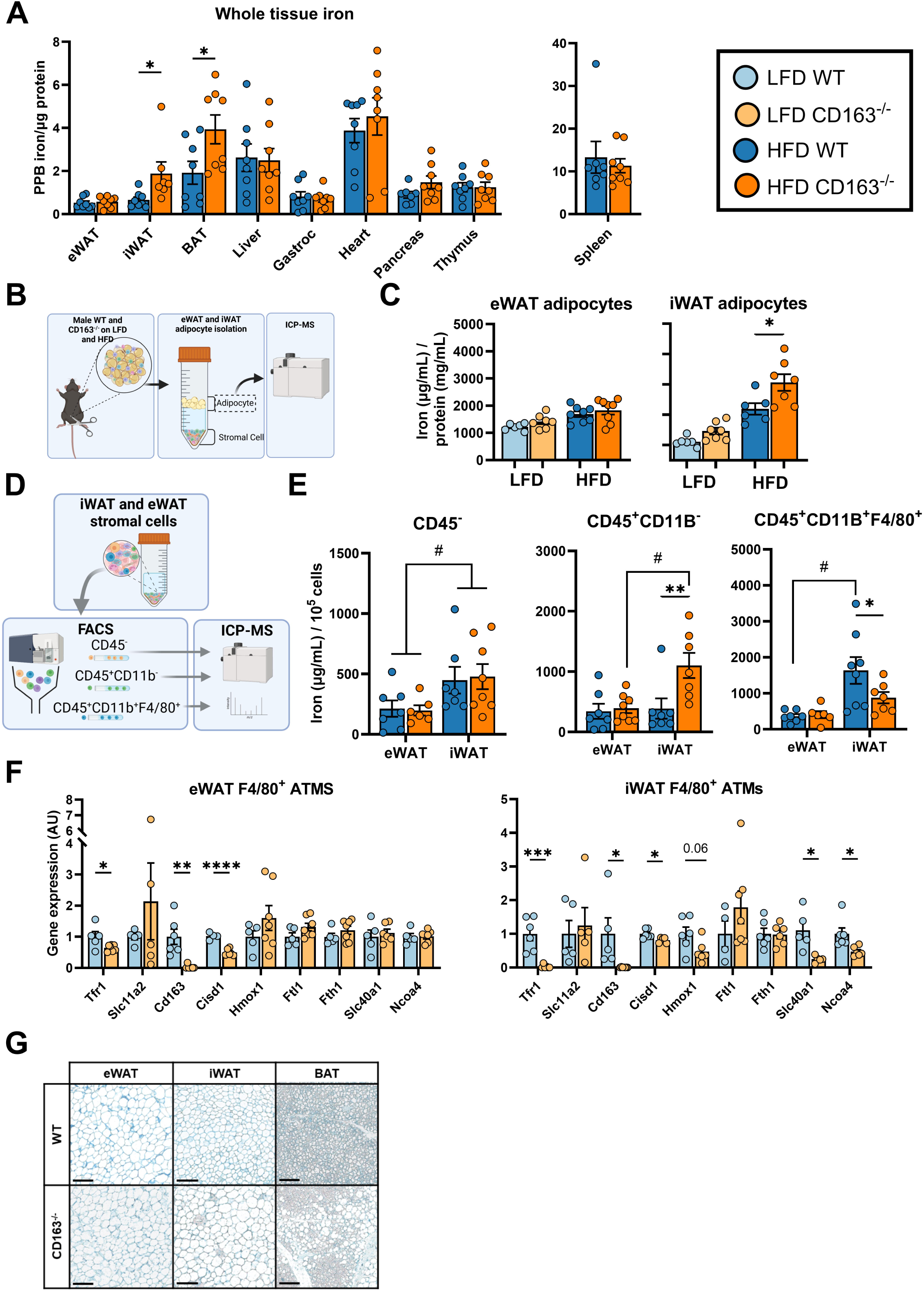
CD163 deficiency causes iWAT iron overload in obesity. A) Whole tissue iron (^56^Fe) measured in eWAT, iWAT, BAT, liver, gastrocnemius skeletal muscle, heart, pancreas, thymus, and spleen. B-C) Adipocytes from eWAT and iWAT were isolated by collagenase digestion (B), and iron within the adipocyte compartment was analyzed by ICP-MS (C). D) Stromal vascular cells from eWAT and iWAT were separated by FACS into CD45^-^ non-hematopoietic cells, CD45^+^CD11B^-^ lymphoid cells, and CD45^+^CD11B^+^F4/80^+^ macrophages. E) Total iron quantified from the CD45^-^, CD45^+^CD11B^-^, and CD45^+^CD11B^+^F4/80^+^ stromal cell populations within eWAT and iWAT. F) Gene signatures related to iron handling (e.g., uptake, storage, export) isolated from F4/80^+^ eWAT and iWAT ATMs. G) Potassium ferrocyanide (Perl’s Prussian blue) staining in eWAT, iWAT, and BAT. Unpaired Student’s *t*-test was used to compare CD163^-/-^ vs. WT littermates for panels A, C, and F. A two-way ANOVA was used to compare iron content with main effects for genotype (Cd163^-/-^ vs. WT) and tissue depot (eWAT vs. iWAT) in panel E. Pairwise comparisons were adjusted with Tukey correction. Data are expressed as mean ± SEM of 6-8 mice per group. ATM, adipose tissue macrophage; BAT, brown adipose tissue; eWAT, epididymal white adipose tissue; FACS, fluorescence-activated cell sorting; Gastroc, gastrocnemius; iWAT, inguinal white adipose tissue; HFD, high-fat diet; ICP-MS, inductively coupled plasma mass spectrometry; LFD, low-fat diet; PPB, parts per billion. *p<0.05, **p<0.01, ***p<0.001, ****p<0.0001; genotype effect, WT vs. Cd163^-/-^. ^#^p<0.05; tissue effect, eWAT vs. iWAT.

### Anti-inflammatory phenotype of CD163^-/-^ BMDMs is diminished upon M2 polarization

To differentiate the impact of CD163 on transcriptomic profile between classically activated (M1-like) and alternatively activated (M2-like) macrophages, BMDMs were isolated and polarized for RNAseq analysis (**Figure 4A**). Principal component analysis was used for dimensional reduction of M1-like and M2-like polarized BMDMs and segregation by genotypes (CD163^-/-^ vs. WT) was found only in M2-like BMDMs (**Figure 4B**). Differential expression analysis in M1-like BMDMs detected 380 upregulated genes (FDR<0.05 & FC>2.0) and 170 significantly downregulated genes in CD163^-/-^ vs. WT BMDMs, whereas M2-like BMDMs presented 351 significantly upregulated genes and 222 downregulated genes in CD163^-/-^ BMDMs (**Figure 4C**). No genes significantly differed between genotypes in M0 BMDMs (**Supplementary Figure 5**). Gene ontology comparing differentially expressed genes between groups identified several redundant pathways significantly altered in M1- and M2-like BMDMs between genotypes. M1-like CD163^-/-^ BMDMs diverged in processes related to ribosomal RNA processing and translation, whereas M2-like CD163^-/-^ BMDMs upregulated several immune and inflammatory processes (**Figure 4D**). Importantly, several M2-like gene signatures (*Arg1*, *Egr2*, *Acadm*, and *Cd200r*) were downregulated in CD163^-/-^ BMDMs upon M2 polarization (**Figure 4E**). Additionally, 42 genes from the innate immune response biological process (GO:0045087) were differentially expressed between genotypes in M2-like BMDMs, all of which were upregulated in CD163^-/-^ compared to WT littermates (**Figure 4F**). Several hemoglobin and iron handling genes from the iron transport biological process (GO:0006826) were also differentially expressed in CD163^-/-^ BMDMs upon M2 polarization (**Figure 4F**).

**Figure 4.**
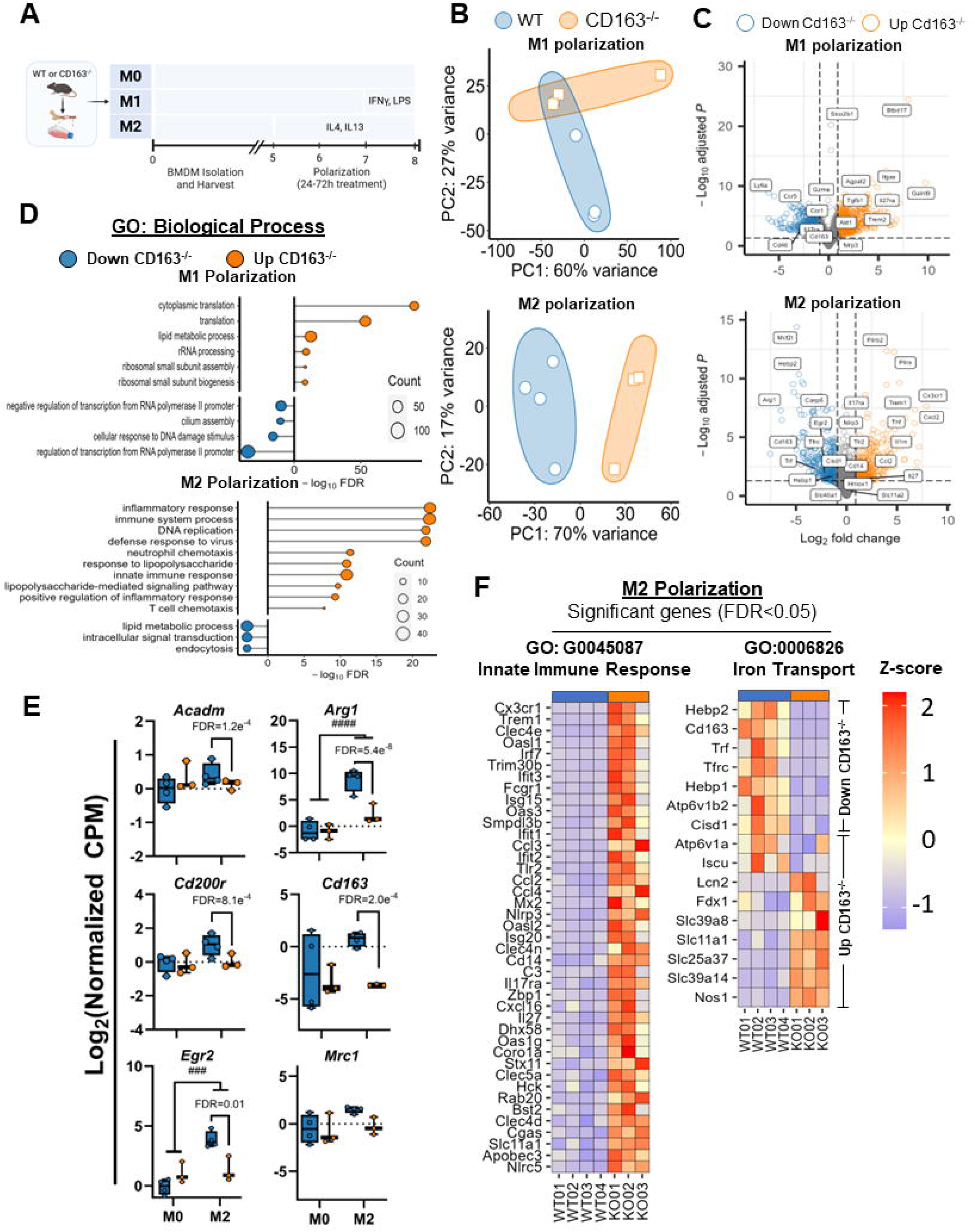
Anti-inflammatory phenotype of CD163^-/-^ BMDMs is diminished upon M2 polarization. A) Bulk RNAseq was performed on BMDMs isolated from CD163^-/-^ and WT littermates (n=3-4/group) and polarized to either M1-like (24h LPS [10 ng/mL] + IFNγ [100 ng/mL]) or M2-like (72h IL-4 [10 ng/mL] + IL-13 [10 ng/mL]) phenotypes. B) Principle component analysis for BMDMs polarized towards M1- and M2-like polarization. C) Inflammatory and iron handling genes from BMDMs polarized towards M1- and M2-like polarization (all FDR<0.05) displayed on volcano plots. D) Gene ontology for significantly upregulated and downregulated biological processes sorted by FDR. E) M2-like polarization gene signatures upon M2 polarization in CD163^-/-^ and WT littermates. F) Differentially expressed Innate Immune Response (GO:0045087) and Iron Transport (GO:0006826) genes (FDR<0.05) from M2-like BMDMs. BMDM, bone marrow derived macrophage; CPM, counts per million; FDR, false discovery rate; GO, gene ontology; IFNγ, interferonLgamma; LPS, lipopolysaccharides; PC, principle component. ### p<0.0001, #### p<0.00001; polarization main effect.

### Loss of CD163 impairs mitochondrial bioenergetics across the BMDM polarization spectrum

Oxygen consumption rate (OCR) and extracellular acidification rate (ECAR) were measured in M0, M1-, and M2-like BMDMs by extracellular flux analysis. Loss of CD163 decreased OCRs with significantly decreased basal, ATP-linked and maximal respiration in M0 BMDMs (**Figure 5A**). M1-like BMDMs from CD163^-/-^ BMDMs presented lower OCR, respiratory measures, and proton-leak (**Figure 5B**). M2-like CD163^-/-^ BMDMs presented lower OCR values including basal, ATP-linked, and maximal respiration (**Figure 5C**). Maximal ECAR during the mitochondrial stress test was greater in M0 and M1-like CD163^-/-^ BMDMs (**Figure 5D**).

**Figure 5.**
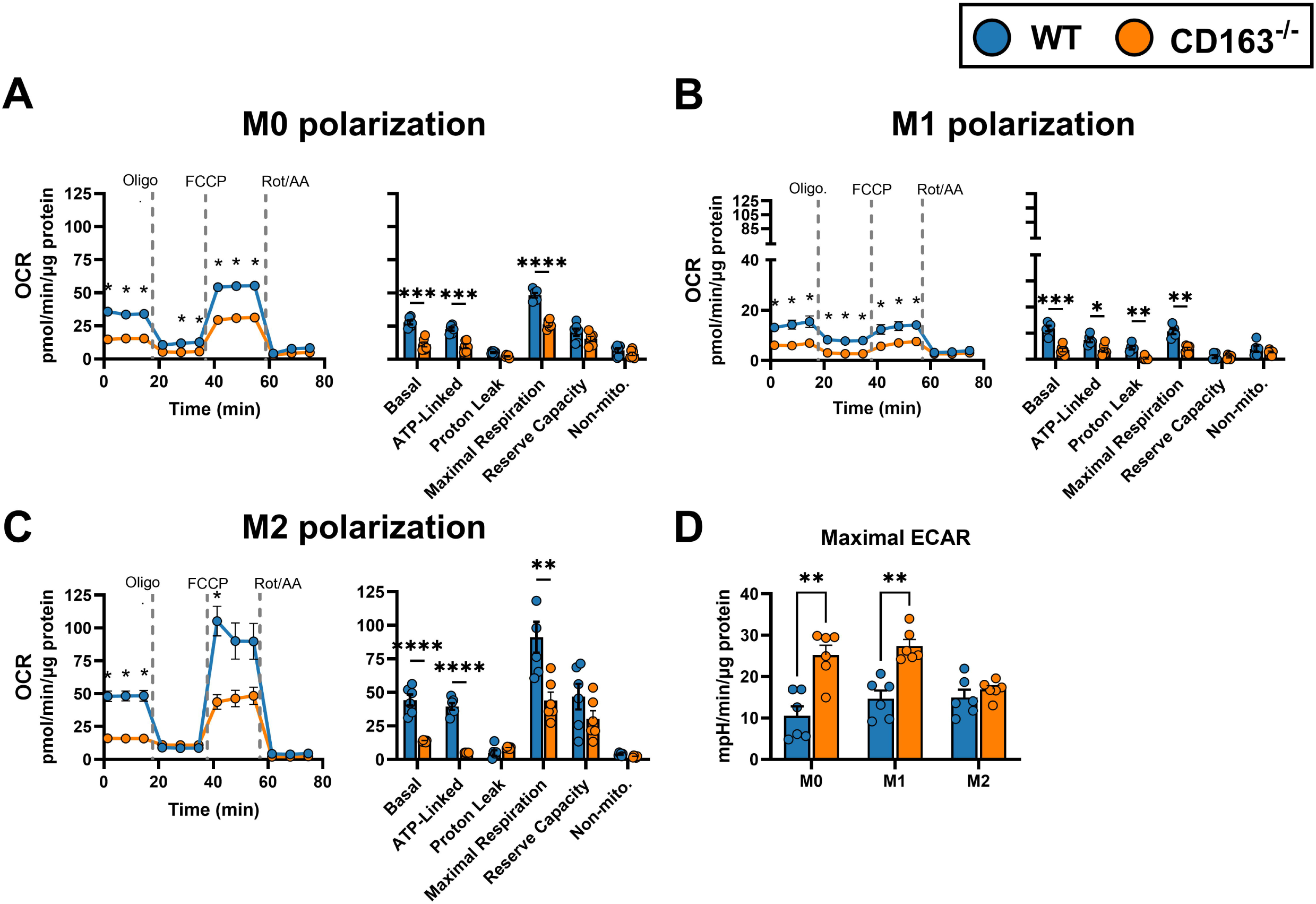
Loss of CD163 impairs mitochondrial bioenergetics across the BMDM polarization spectrum. BMDMs were isolated from CD163^-/-^ and WT littermates then polarized to either M1-like (24h LPS [10 ng/mL] + IFNγ [100 ng/mL]) or M2-like (72h IL-4 [10 ng/mL] + IL-13[10 ng/mL]) phenotypes, and extracellular flux analyses were conducted using the Seahorse Analyzer. A-C) OCR curves and OCR parameters including basal respiration, ATP-linked respiration, proton leak, maximal respiratory capacity, reserve capacity, and non-mitochondrial oxidation in M0 (A), M1-(B), and M2-like BMDMs (C). D) Maximal ECAR measured during the mitochondrial stress test. Unpaired Student’s *t*-test was used to compare CD163^-/-^ vs. WT littermates during the mitochondrial stress test and OCR parameters for panels B-E. Data are expressed as mean ± SEM of 4-6 mice/group. ECAR, extracellular acidification rate; FCCP, Carbonyl cyanide 4-(trifluoromethoxy)phenylhydrazone; IFNγ, interferonLgamma, LPS, lipopolysaccharides; Oligo., oligomycin; OCR, oxygen consumption rate; Rot/AA, rotenone & antimycin A. *p<0.05, **p<0.01, ***p<0.001, ****p<0.0001; genotype affect, WT vs. Cd163^-/-^.

### Loss of CD163 reduces hemoglobin uptake by M2-like macrophages

Hemoglobin uptake was quantified in M0, M1-, and M2-like BMDMs treated with 25 mg/dL fluorescence conjugated hemoglobin for 24 h (**Figure 6A and Supplementary Figure 6**). Flow cytometry was used to measure median fluorescence intensity (MFI) as a proxy for hemoglobin uptake, in BMDMs treated with hemoglobin for 2, 6, and 24 h (**Figure 6B**). Over the 24 h treatment period, M2-like BMDMs presented >2-fold greater hemoglobin uptake compared to M0 and M1-like BMDMs (**Figure 6C**). Further, in M2-like BMDMs, CD163 deficiency attenuated hemoglobin uptake after 24 h (**Figure 6C**). Total hemoglobin-positive area was also measured by fluorescence imaging (**Figure 6D**). M0 BMDMs presented greater hemoglobin-positive area at 6 h, and returned to near basal levels after 24 h, whereas M1-like BMDMs increased after 24 h, and was greater in WT littermates (**Figure 6E**). M2-like BMDMs significantly increased hemoglobin-postive area at 6 h and throughout the 24 h treatment only in WT BMDMs, whereas CD163^-/-^ BMDMs presented area fractions similar to M0 and M1-like BMDMs (**Figure 6E**). These data show loss of CD163 hinders M2-like macrophage potential to buffer extracellular free hemoglobin.

**Figure 6.**
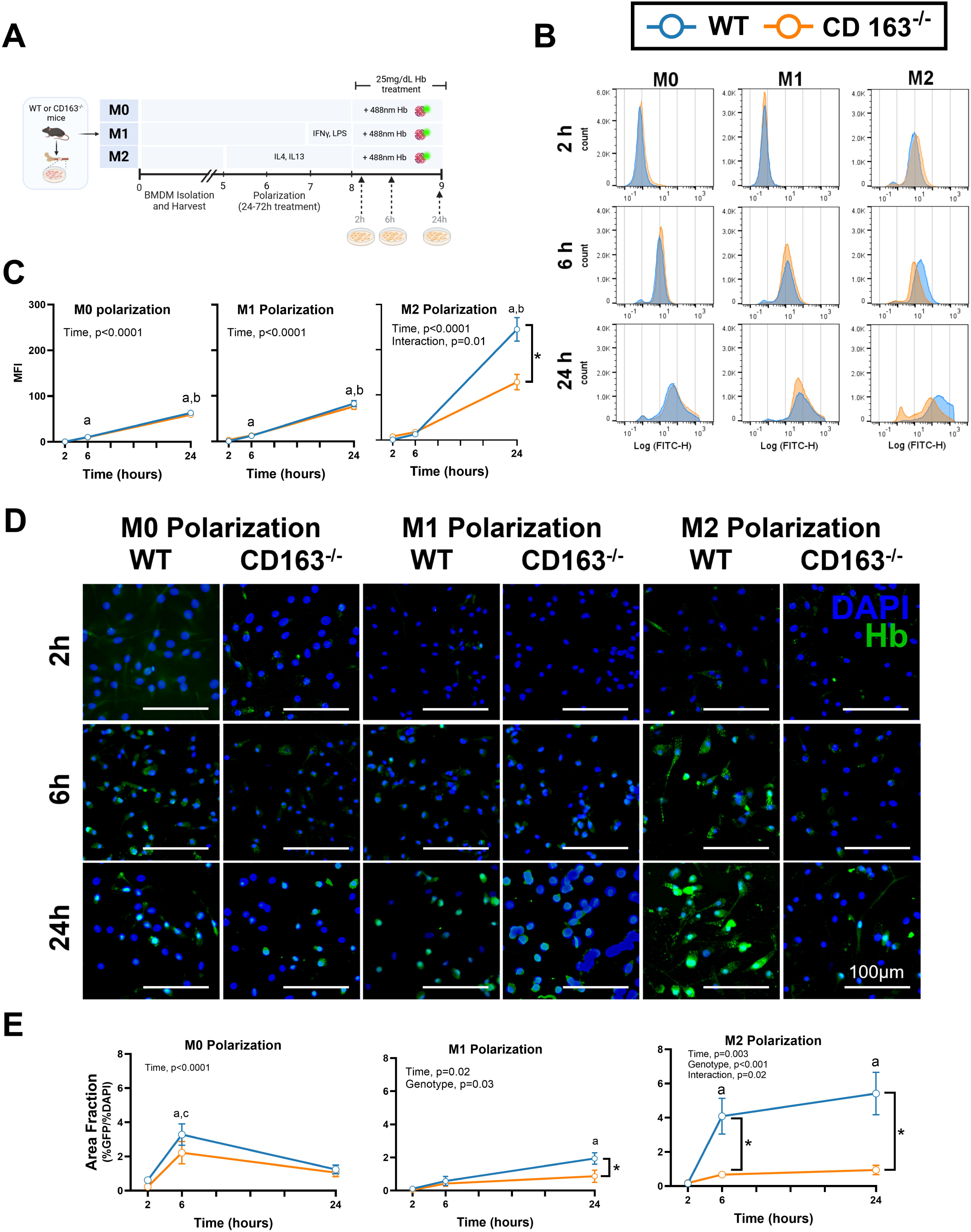
Loss of CD163 reduces hemoglobin uptake by M2-like macrophages. A) BMDMs were isolated from CD163^-/-^ and WT littermates, then polarized to either M1-like (24h LPS [10ng/mL] + IFNγ [100ng/mL]) or M2-like (72h IL-4 [10ng/mL] + IL-13 [10ng/mL]) phenotypes, then treated with 25 mg/dL Hb conjugated to 488nm fluorophore for 2, 6, and 24 h. B) Representative MFI histogram comparing Hb-treated M0, M1-, and M2-like BMDMs in CD163^-/-^ vs. WT littermates for 2,6, and 24 h. C) Mean MFI from 4 biological replicates per group for each Hb treatment time point. D) Representative immunofluorescence images for BMDMs labeled with Hb-conjugated fluorophore (488nm; green) and DAPI (blue). Images collected at 40X magnification and scale bar specifies 100μm. E) Area fraction measured by % Hb^+^ area normalized to % DAPI^+^ area for polarized BMDMs treated for 2, 6, and 24 h of 3-4 mice per group. A two-way ANOVA compared MFI and area fraction with main effects including genotype (WT vs. CD163^-/-^) and treatment time (2, 6, 24 h) for M0, M1-, and M2-like BMDMs in panel C and E. BMDMs, bone marrow derived macrophages; Hb, hemoglobin; IFNγ, interferonLgamma; LPS, lipopolysaccharides; MFI, median fluorescence intensity. Data are expressed as mean ± SEM of 3-4 mice per polarization group and time point. *p<0.05; genotype main effect, WT vs. Cd163^-/-^; a-c, time main effect, a, p<0.05 vs. 2h; b, p<0.05 vs. 6h; c, p<0.05 vs. 24h.

## Discussion

Iron is a critical element for numerous biological processes, and is exquisitely controlled by tissue- and cellular-dependent processes (2). However, excess iron is harmful to cells and is associated with β cell dysfunction, nonalcoholic fatty liver diseas, and type 2 diabetes risk (1). Obesity results in pro-inflammatory macrophage recruitment into adipose tissue (13-15), and macrophage polarization is tightly linked with iron handling (9, 30). Our research group identified a subgroup of tissue-resident macrophages, MFe^hi^ cells, with high intrinsic capacity for iron cycling (17, 31). These cells are marked with greater CD163 and tissue-resident macrophage gene expression, suggesting the anti-inflammatory influence of this receptor is partly explained by free hemoglobin scavenging. However, the capacity for iron handling in MFe^hi^ cells becomes diminished upon obesity (17). Here, we show that CD163 is critical to support an anti-inflammatory phenotype in M2-like BMDMs, suggesting tissue-resident macrophages *in vivo* may become compromised upon CD163 deficiency. These findings support CD163 as a critical receptor towards limiting severe insulin resistance upon obesity, partially through its anti-inflammatory contribution to free hemoglobin scavenging.

Hemolysis is a physiological process by which hemoglobin is released into the circulation from senescent erythrocytes. Free hemoglobin has a strong affinity for haptoglobin, primarily produced by hepatocytes, protecting tissues from heme-induced oxidative damage (32, 33). However, the antioxidant potential for haptoglobin requires CD163 on tissue-resident macrophages to scavenge Hb-Hp complexes. Additionally, heme catabalism by heme-oxygenase I results in the production of free iron, carbon monoxide, and biliverdin; and CO production is noted to have anti-inflammatory effects by IL-10 release (34). Others have shown that excess cellular heme-iron initiates the release of pro-inflammatory cytokines from macrophages (35-37). Therefore, heme catabolism (and lower circulating free heme) appears vital to sustain anti-inflammatory roles of CD163-expressing macrophages. Here, we confirmed M2-like macrophages have a profound role for hemoglobin uptake compared to M0 and M1-like BMDMs, and loss of CD163 nearly abolishes the hemoglobin-regulatory components signature of M2-like macrophages. These data support a critical role for CD163 in buffering iron-containing heme, and limiting iron spillover into key metabolic tissues.

Macrophage metabolic states are linked to their polarization (38). Specifically, M1-like macrophages primarily rely on glycolysis, while M2-like macrophages are oxidative (39). Previous reports have shown both mitochondrial iron overload and depletion impair oxidative metabolism (19). We observed loss of CD163 attenuated mitochondrial oxidation across the BMDM polarization spectra, and M2-like macrophages had the greatest absolute OCR impairment. It is unknown whether OCR impairments in complexes I, II, and III are due to perturbations in Fe-S cluster regulation. *Cisd1 (*MitoNEET), an inhibitor of mitochondrial iron uptake, was decreased in CD163^-/-^ eWAT and iWAT ATMs. Recent reports have shown macrophage-specific MitoNEET overexpression lowers mitochondrial matrix iron content and promotes an anti-inflammatory ATM profile, protecting from insulin resistance (19). We speculate the lower heme- and transferrin-derived iron uptake in CD163^-/-^ BMDMs lessens cytosolic iron accumulation, but increases mitochondrial iron accumulation as shown by decreased MitoNEET expression in ATMs. This effect may explain why M2-like BMDMs upregulate inflammatory gene signatures, because lipid uptake is utilized as a pro-inflammatory mediator in M1-like macrophages (e.g., eicosanoids), as opposed to oxidation in M2-like macrophages (40). Whether macrophage mitochondrial metabolism dictates the fate of polarization, or vice versa, and the influence of iron in this process, will be intriguing to uncover in future work.

It is established that macrophage inflammatory activation is related to iron-handling capacity (9). Recent reports have shown depot-specific differences in inflammatory cell profile, where iWAT present ∼70% higher proportion of M2-like ATMs compared to eWAT (41). We found loss of CD163 resulted in a depot-specific differences in iron deposition in BAT and iWAT adipocytes upon HFD. Increased iWAT adipocyte iron was accompanied by lower iWAT ATM iron, suggesting CD163 deficiency impedes the iron buffering capacity of iWAT ATMs, resulting in iron spillover into adipocytes and other immune cells. Additionally, iWAT iron overload suggests that metabolic dysfunction upon HFD may be temporally initiated in this ‘subcutaneous-like’ adipose depot, and follows with systemic consequences to glucose tolerance. In humans, the majority of lipolysis-derived FA originate from subcutaneous adipose tissue compared to visceral sources [e.g., eWAT; (42)]. Loss of CD163 also attenuated *Tfr1* expression in M2-like BMDMs and eWAT/iWAT ATMs, which is consistent with previous reports showing M2-like macrophages present higher levels of *Tfr1* and *Cd163* in the prescence of supraphysiological iron concentration in effort to buffer extracellular iron and/or support tissue repair processes (43). Therefore, the pro-inflammatory landscape of CD163 deficient ATMs may also impair iron homeostatic functions by limiting Tf-bound iron uptake, in addition to the observed differences in Hb-Hp receptor scavenging.

The cross-talk linking macrophage and adipocyte iron regulation with systemic insulin sensitivity are still unconfirmed. Here, iWAT adipocytes were burdened with greater iron, likely consequential to diminished ATM iron clearance. Further, NEFA content was elevated during the clamp period of the hyperinsulinemic-euglycempic clamp, which is a consequence of greater postprandial lipolysis and lower FA clearance. In humans, the subcutaneous adipose tissue corresponds to ∼70% of total lipolysis compared to visceral sources (42), and greater lipolysis rates upon hyperinsulinemia are linked with poor insulin-mediated glucose disposal (44). Because skeletal muscle glucose uptake was impaired during the hyperinsulemic clamp, and skeletal muscle is the primary source for insulin-mediated glucose disposal (45), we propose adipocyte iron overload accelerates postprandial lipolysis in accordance with skeletal muscle FA uptake. Further, we show a pro-inflammatory phenotype in M2-like BMDMs may also translate to greater pro-inflamatory cytokine release by ATMs (e.g., IL6, TNFα, IL-1β); a hallmark of obesity-induced adipose tissue inflammation (12). These cytokines may also also sustain a cycle of pro-inflammatory immune cell infiltration and local insulin resistance in adipose tissue (46), resulting in excess lipolysis and FA intermediate production known impair insulin action.

We report that CD163 deficiency results in immunometabolic defects including reduced mitochondrial metabolism, loss of anti-inflammatory phenotype in M2-like BMDMs, and impaired hemoglobin scavenging. To this end, systemic insulin resistance was present, in addition to iWAT adipocyte iron overload. Future studies will delineate whether loss of CD163 induces insulin resistance as a consequence of iron overload in metabolic tissues or by loss of anti-inflammatory function; both roles are key signatures for CD163 in tissue-resident macrophage.

## Acknowledgements

We acknowledge Marnie Gruen for assistance with animal husbandry, and Gabriel Ferguson for assistance with experiments. We thank Stuart Landstreet and Dr. Ciara Shaver for assistance with fluorescence hemoglobin preparation and guidance. We acknowledge the following Vanderbilt University (VU) and Vanderbilt University Medical Center (VUMC) core facilities: VUMC Hormone Assay & Analytical Services Core (NIH DK020593), VU Metabolic Mouse Phenotyping Center [VMMPC (NIH DK059637; www.vmmpc.org)], Translational Pathology Shared Resource (NCI/NIH Cancer Center Support Grant 5P30 CA68485-19), Vanderbilt University Medical Center Flow Cytometry Shared Resource Core (DK058404), and Vanderbilt Mass Spectrometry Research Center.

## Data and Resource Availability

RNAseq data and source code used fro this publication are accessible through Github data repository. https://github.com/MichaelSchleh/CD163-and-immunometabolic-health

## Author Contributions

M.W.S. designed research studies, performed experiments, interpreted results, and drafted the manuscript. A.R. was responsible for mouse phenotyping, animal husbandry, performed experiments, and interpreted results. M.A. conceived and design research studies, and edited the manuscript. M.A. M.A. currently holds a position with the University of Nairobi, Department of Medical Microbiology and Immunology. A.H.H. conceived and designed research studies, interpreted results, and reviewed and edited the manuscript. All authors approved the final version of the manuscript. A.H.H. is the guarantor of this work and, as such, had full access to all the data in the study and takes responsibility for the integrity of the data and the accuracy of the data analysis.

## References

1. Harrison AV, Lorenzo FR, McClain DA. Iron and the Pathophysiology of Diabetes. Annual Review of Physiology. 2023;85(1):339–62. doi: 10.1146/annurev-physiol-022522-102832. PubMed PMID: 36137277.

2. Galy B, Conrad M, Muckenthaler M. Mechanisms controlling cellular and systemic iron homeostasis. Nature Reviews Molecular Cell Biology. 2023. doi: 10.1038/s41580-023-00648-1.

3. Kautz L, Nemeth E. Molecular liaisons between erythropoiesis and iron metabolism. Blood. 2014;124(4):479–82. Epub 2014/05/31. doi: 10.1182/blood-2014-05-516252. PubMed PMID: 24876565; PMCID: PMC4110655.

4. Zhao J, Andreev I, Silva HM. Resident tissue macrophages: Key coordinators of tissue homeostasis beyond immunity. Science Immunology. 2024;9(94):eadd1967. doi: doi:10.1126/sciimmunol.add1967.

5. Winn NC, Volk KM, Hasty AH. Regulation of tissue iron homeostasis: the macrophage "ferrostat". JCI Insight. 2020;5(2). Epub 2020/01/31. doi: 10.1172/jci.insight.132964. PubMed PMID: 31996481; PMCID: PMC7098718 exists.

6. Ginhoux F, Guilliams M. Tissue-Resident Macrophage Ontogeny and Homeostasis. Immunity. 2016;44(3):439-49. Epub 2016/03/18. doi: 10.1016/j.immuni.2016.02.024. PubMed PMID: 26982352.

7. Coats BR, Schoenfelt KQ, Barbosa-Lorenzi VC, Peris E, Cui C, Hoffman A, Zhou G, Fernandez S, Zhai L, Hall BA, Haka AS, Shah AM, Reardon CA, Brady MJ, Rhodes CJ, Maxfield FR, Becker L. Metabolically Activated Adipose Tissue Macrophages Perform Detrimental and Beneficial Functions during Diet-Induced Obesity. Cell Rep. 2017;20(13):3149–61. Epub 2017/09/28. doi: 10.1016/j.celrep.2017.08.096. PubMed PMID: 28954231; PMCID: PMC5646237.

8. Kratz M, Coats BR, Hisert KB, Hagman D, Mutskov V, Peris E, Schoenfelt KQ, Kuzma JN, Larson I, Billing PS, Landerholm RW, Crouthamel M, Gozal D, Hwang S, Singh PK, Becker L. Metabolic dysfunction drives a mechanistically distinct proinflammatory phenotype in adipose tissue macrophages. Cell Metab. 2014;20(4):614–25. Epub 2014/09/23. doi: 10.1016/j.cmet.2014.08.010. PubMed PMID: 25242226; PMCID: PMC4192131.

9. Gianfranca C, Lara C, Emanuele P, Alessandra C, Enrico T, Lidia B, Alessandro C, Silvia B, Angelo AM, Pietro A, Laura S, Clara C, Patrizia R-Q. Polarization dictates iron handling by inflammatory and alternatively activated macrophages. Haematologica. 2010;95(11):1814–22. doi: 10.3324/haematol.2010.023879.

10. Kristiansen M, Graversen JH, Jacobsen C, Sonne O, Hoffman HJ, Law SK, Moestrup SK. Identification of the haemoglobin scavenger receptor. Nature. 2001;409(6817):198-201. Epub 2001/02/24. doi: 10.1038/35051594. PubMed PMID: 11196644.

11. Jacks RD, Lumeng CN. Macrophage and T cell networks in adipose tissue. Nature Reviews Endocrinology. 2024;20(1):50–61. doi: 10.1038/s41574-023-00908-2.

12. Schleh MW, Caslin HL, Garcia JN, Mashayekhi M, Srivastava G, Bradley AB, Hasty AH. Metaflammation in obesity and its therapeutic targeting. Science Translational Medicine. 2023;15(723):eadf9382. doi: doi:10.1126/scitranslmed.adf9382.

13. Weisberg SP, McCann D, Desai M, Rosenbaum M, Leibel RL, Ferrante AW, Jr. Obesity is associated with macrophage accumulation in adipose tissue. J Clin Invest. 2003;112(12):1796–808. Epub 2003/12/18. doi: 10.1172/jci19246. PubMed PMID: 14679176; PMCID: PMC296995.

14. Lumeng CN, Bodzin JL, Saltiel AR. Obesity induces a phenotypic switch in adipose tissue macrophage polarization. The Journal of Clinical Investigation. 2007;117(1):175–84. doi: 10.1172/JCI29881.

15. Xu H, Barnes GT, Yang Q, Tan G, Yang D, Chou CJ, Sole J, Nichols A, Ross JS, Tartaglia LA, Chen H. Chronic inflammation in fat plays a crucial role in the development of obesity-related insulin resistance. J Clin Invest. 2003;112(12):1821–30. Epub 2003/12/18. doi: 10.1172/jci19451. PubMed PMID: 14679177; PMCID: PMC296998.

16. Saltiel AR, Olefsky JM. Inflammatory mechanisms linking obesity and metabolic disease. The Journal of Clinical Investigation. 2017;127(1):1–4. doi: 10.1172/JCI92035.

17. Orr JS, Kennedy A, Anderson-Baucum EK, Webb CD, Fordahl SC, Erikson KM, Zhang Y, Etzerodt A, Moestrup SK, Hasty AH. Obesity alters adipose tissue macrophage iron content and tissue iron distribution. Diabetes. 2014;63(2):421–32. Epub 2013/10/17. doi: 10.2337/db13-0213. PubMed PMID: 24130337; PMCID: PMC3900546.

18. Ameka MK, Beavers WN, Shaver CM, Ware LB, Kerchberger VE, Schoenfelt KQ, Sun L, Koyama T, Skaar EP, Becker L, Hasty AH. An Iron Refractory Phenotype in Obese Adipose Tissue Macrophages Leads to Adipocyte Iron Overload. Int J Mol Sci. 2022;23(13). Epub 2022/07/10. doi: 10.3390/ijms23137417. PubMed PMID: 35806422; PMCID: PMC9267114.

19. Joffin N, Gliniak CM, Funcke JB, Paschoal VA, Crewe C, Chen S, Gordillo R, Kusminski CM, Oh DY, Geldenhuys WJ, Scherer PE. Adipose tissue macrophages exert systemic metabolic control by manipulating local iron concentrations. Nat Metab. 2022;4(11):1474–94. Epub 2022/11/05. doi: 10.1038/s42255-022-00664-z. PubMed PMID: 36329217.

20. Tang Y, Wang D, Zhang H, Zhang Y, Wang J, Qi R, Yang J, Shen H, Xu Y, Li M. Rapid responses of adipocytes to iron overload increase serum TG level by decreasing adiponectin. J Cell Physiol. 2021;236(11):7544–53. Epub 2021/04/16. doi: 10.1002/jcp.30391. PubMed PMID: 33855731.

21. Dongiovanni P, Ruscica M, Rametta R, Recalcati S, Steffani L, Gatti S, Girelli D, Cairo G, Magni P, Fargion S, Valenti L. Dietary iron overload induces visceral adipose tissue insulin resistance. Am J Pathol. 2013;182(6):2254–63. Epub 2013/04/13. doi: 10.1016/j.ajpath.2013.02.019. PubMed PMID: 23578384.

22. Gabrielsen JS, Gao Y, Simcox JA, Huang J, Thorup D, Jones D, Cooksey RC, Gabrielsen D, Adams TD, Hunt SC, Hopkins PN, Cefalu WT, McClain DA. Adipocyte iron regulates adiponectin and insulin sensitivity. J Clin Invest. 2012;122(10):3529–40. Epub 2012/09/22. doi: 10.1172/jci44421. PubMed PMID: 22996660; PMCID: PMC3461897.

23. Cottam MA, Caslin HL, Winn NC, Hasty AH. Multiomics reveals persistence of obesity-associated immune cell phenotypes in adipose tissue during weight loss and weight regain in mice. Nature Communications. 2022;13(1):2950. doi: 10.1038/s41467-022-30646-4.

24. Orr JS, Kennedy AJ, Hasty AH. Isolation of adipose tissue immune cells. J Vis Exp. 2013(75):e50707. Epub 2013/06/04. doi: 10.3791/50707. PubMed PMID: 23728515; PMCID: PMC3718226.

25. Love MI, Huber W, Anders S. Moderated estimation of fold change and dispersion for RNA-seq data with DESeq2. Genome Biology. 2014;15(12):550. doi: 10.1186/s13059-014-0550-8.

26. Volk Robertson K, Schleh MW, Harrison FE, Hasty AH. Microglial-specific knockdown of iron import gene, Slc11a2, blunts LPS-induced neuroinflammatory responses in a sex-specific manner. Brain, Behavior, and Immunity. 2024;116:370-84. doi: 10.1016/j.bbi.2023.12.020.

27. Benjamini Y, Hochberg Y. Controlling the False Discovery Rate: A Practical and Powerful Approach to Multiple Testing. Journal of the Royal Statistical Society Series B (Methodological). 1995;57(1):289–300.

28. Winn NC, Wolf EM, Cottam MA, Bhanot M, Hasty AH. Myeloid-specific deletion of ferroportin impairs macrophage bioenergetics but is disconnected from systemic insulin action in adult mice. American Journal of Physiology-Endocrinology and Metabolism. 2021;321(3):E376–E91. doi: 10.1152/ajpendo.00116.2021.

29. Ascaso JF, Pardo S, Real JT, Lorente RI, Priego A, Carmena R. Diagnosing insulin resistance by simple quantitative methods in subjects with normal glucose metabolism. Diabetes Care. 2003;26(12):3320–5. Epub 2003/11/25. doi: 10.2337/diacare.26.12.3320. PubMed PMID: 14633821.

30. Recalcati S, Locati M, Marini A, Santambrogio P, Zaninotto F, De Pizzol M, Zammataro L, Girelli D, Cairo G. Differential regulation of iron homeostasis during human macrophage polarized activation. Eur J Immunol. 2010;40(3):824–35. Epub 2009/12/30. doi: 10.1002/eji.200939889. PubMed PMID: 20039303.

31. Hubler MJ, Erikson KM, Kennedy AJ, Hasty AH. MFe(hi) adipose tissue macrophages compensate for tissue iron perturbations in mice. Am J Physiol Cell Physiol. 2018;315(3):C319-c29. Epub 2018/05/17. doi: 10.1152/ajpcell.00103.2018. PubMed PMID: 29768045; PMCID: PMC6171041.

32. Melamed-Frank M, Lache O, Enav BI, Szafranek T, Levy NS, Ricklis RM, Levy AP. Structure-function analysis of the antioxidant properties of haptoglobin. Blood. 2001;98(13):3693–8. Epub 2001/12/12. doi: 10.1182/blood.v98.13.3693. PubMed PMID: 11739174.

33. McCormick DJ, Atassi MZ. Hemoglobin binding with haptoglobin: delineation of the haptoglobin binding site on the alpha-chain of human hemoglobin. J Protein Chem. 1990;9(6):735–42. Epub 1990/12/01. doi: 10.1007/bf01024768. PubMed PMID: 2073325.

34. Ryter SW, Choi AM. Targeting heme oxygenase-1 and carbon monoxide for therapeutic modulation of inflammation. Transl Res. 2016;167(1):7–34. Epub 2015/07/15. doi: 10.1016/j.trsl.2015.06.011. PubMed PMID: 26166253; PMCID: PMC4857893.

35. Vinchi F, Costa da Silva M, Ingoglia G, Petrillo S, Brinkman N, Zuercher A, Cerwenka A, Tolosano E, Muckenthaler MU. Hemopexin therapy reverts heme-induced proinflammatory phenotypic switching of macrophages in a mouse model of sickle cell disease. Blood. 2016;127(4):473–86. Epub 2015/12/18. doi: 10.1182/blood-2015-08-663245. PubMed PMID: 26675351; PMCID: PMC4850229.

36. Fortes GB, Alves LS, de Oliveira R, Dutra FF, Rodrigues D, Fernandez PL, Souto-Padron T, De Rosa MJ, Kelliher M, Golenbock D, Chan FKM, Bozza MT. Heme induces programmed necrosis on macrophages through autocrine TNF and ROS production. Blood. 2012;119(10):2368–75. doi: 10.1182/blood-2011-08-375303.

37. Figueiredo RT, Fernandez PL, Mourao-Sa DS, Porto BN, Dutra FF, Alves LS, Oliveira MF, Oliveira PL, Graça-Souza AV, Bozza MT. Characterization of Heme as Activator of Toll-like Receptor 4*. Journal of Biological Chemistry. 2007;282(28):20221–9. doi: 10.1074/jbc.M610737200.

38. Caslin HL, Hasty AH. Extrinsic and Intrinsic Immunometabolism Converge: Perspectives on Future Research and Therapeutic Development for Obesity. Curr Obes Rep. 2019;8(3):210–9. Epub 2019/03/29. doi: 10.1007/s13679-019-00344-2. PubMed PMID: 30919312; PMCID: PMC6661206.

39. Rodríguez-Prados JC, Través PG, Cuenca J, Rico D, Aragonés J, Martín-Sanz P, Cascante M, Boscá L. Substrate fate in activated macrophages: a comparison between innate, classic, and alternative activation. J Immunol. 2010;185(1):605–14. Epub 2010/05/26. doi: 10.4049/jimmunol.0901698. PubMed PMID: 20498354.

40. Remmerie A, Scott CL. Macrophages and lipid metabolism. Cell Immunol. 2018;330:27-42. Epub 2018/02/13. doi: 10.1016/j.cellimm.2018.01.020. PubMed PMID: 29429624; PMCID: PMC6108423.

41. Rohm TV, Castellani Gomes Dos Reis F, Isaac R, Murphy C, Cunha e Rocha K, Bandyopadhyay G, Gao H, Libster AM, Zapata RC, Lee YS, Ying W, Miciano C, Wang A, Olefsky JM. Adipose tissue macrophages secrete small extracellular vesicles that mediate rosiglitazone-induced insulin sensitization. Nature Metabolism. 2024. doi: 10.1038/s42255-024-01023-w.

42. Nielsen S, Guo Z, Johnson CM, Hensrud DD, Jensen MD. Splanchnic lipolysis in human obesity. The Journal of Clinical Investigation. 2004;113(11):1582–8. doi: 10.1172/JCI21047.

43. Corna G, Campana L, Pignatti E, Castiglioni A, Tagliafico E, Bosurgi L, Campanella A, Brunelli S, Manfredi AA, Apostoli P, Silvestri L, Camaschella C, Rovere-Querini P. Polarization dictates iron handling by inflammatory and alternatively activated macrophages. Haematologica. 2010;95(11):1814–22. Epub 2010/06/01. doi: 10.3324/haematol.2010.023879. PubMed PMID: 20511666; PMCID: PMC2966902.

44. Schleh MW, Ryan BJ, Ahn C, Ludzki AC, Varshney P, Gillen JB, Van Pelt DW, Pitchford LM, Howton SM, Rode T, Chenevert TL, Hummel SL, Burant CF, Horowitz JF. Metabolic dysfunction in obesity is related to impaired suppression of fatty acid release from adipose tissue by insulin. Obesity. 2023;31(5):1347–61. doi: 10.1002/oby.23734.

45. DeFronzo RA, Jacot E, Jequier E, Maeder E, Wahren J, Felber JP. The Effect of Insulin on the Disposal of Intravenous Glucose: Results from Indirect Calorimetry and Hepatic and Femoral Venous Catheterization. Diabetes. 1981;30(12):1000–7. doi: 10.2337/diab.30.12.1000.

46. Hotamisligil GS, Shargill NS, Spiegelman BM. Adipose expression of tumor necrosis factor-alpha: direct role in obesity-linked insulin resistance. Science. 1993;259(5091):87-91. Epub 1993/01/01. doi: 10.1126/science.7678183. PubMed PMID: 7678183.

47. Karlsson M, Zhang C, Méar L, Zhong W, Digre A, Katona B, Sjöstedt E, Butler L, Odeberg J, Dusart P, Edfors F, Oksvold P, von Feilitzen K, Zwahlen M, Arif M, Altay O, Li X, Ozcan M, Mardinoglu A, Fagerberg L, Mulder J, Luo Y, Ponten F, Uhlén M, Lindskog C. A single–cell type transcriptomics map of human tissues. Science Advances. 2021;7(31):eabh2169. doi: doi:10.1126/sciadv.abh2169.

