## Supplementary material for "Deficiency of the hemoglobin-haptoglobin receptor, CD163, worsens insulin sensitivity in obese male mice": Supplementary Information.docx

**Supplementary Methods**

***Hyperinsulinemic-euglycemic Clamp***

One week before the clamp procedure, arterial and venous catheters were surgically inserted in the carotid artery and jugular veins for continuous sampling and infusions respectively. Ninety minutes prior to the clamp (t=-90 min), [3-^3^H]glucose was continuously infused (0.8 µCi prime followed by 0.04 μCi/min continuous infusion). Insulin was continuously infused at t=0 min (4 mU/kg/min), and glucose was monitored every 10 min to adjust the variable glucose infusion rate (GIR) to achieve euglycemia at t=155 min. Infused glucose contained trace amounts (0.06 µCi/µL) of [3-^3^H]glucose. Heparinized erythrocytes from donor mice were infused at 5 µL/min to compensate for blood loss. Clamp insulin was collected at t=100 and t=120 min. At 120 min, 13 μCi of 2[^14^C]deoxyglucose ([^14^C]2DG) was administered to determine tissue-specific glucose uptake (Rg), in which blood samples were collected at 122, 125, 130, 135, and 145 min to measure glucose rate of disappearance (Rd) from plasma. At 145 min, mice were euthanized and tissues immediately flash frozen. Glucose rate of appearance (Ra) and Rd were determined under steady state conditions (1).

***Adipocyte and stromal cell isolation***

The isolation of epididymal white adipose tissue (eWAT) and iWAT adipocytes from the stromal vascular fraction (SVF) was completed by mincing WAT in PBS containing 1% FBS (PBS/FBS), and digesting in 2 mg/mL collagenase type IV (Worthington) for 30 min. For most experiments, left and right side eWAT and iWAT were pooled, and minced samples incubated for 30 min and 60 min for eWAT and iWAT, respectively, while shaking at 200 rpm at 37°C. Samples were then diluted in PBS/FBS, filtered through a 100 μm cell strainer, and centrifuged for 5 min at 500 g at 4°C to pellet the SVF. Floated adipocytes were collected while the SVF pellet was resuspended in red blood cell lysis buffer, and the lysis reaction was quenched by the addition of 10 mL PBS/FBS. The SVF was filtered through a 40 μm cell strainer and centrifuged for 5 min at 500 g at 4°C, where the pelleted cells were analyzed and isolated by flow cytometry, or F4/80^+^ magnetic bead separation (Miltenyi).

***Inductively coupled plasma mass spectrometry (ICP-MS)***

Tissues, adipocytes, and BMDMs were homogenized and digested in 200 μL of Optima-grade nitric acid (VWR) plus 50 μL of Ultratrace-grade hydrogen peroxide (Thermofisher) in metal-free 15 mL conical tubes (VWR) and, then incubated overnight at 65°C. The following day, 2 mL of ultrapure-grade water was added to neutralize the digestion. Total iron from the acid digested samples was quantified by ICP-MS (Agilent 7700, Santa Clara, CA) attached to an autosampler (Teledyne CETAC Technologies, Omaha, NE) at the Vanderbilt Mass Spectrometry Research Center. For analysis, the following settings were used: Cell entrance = -40V, cell exit = -60V, plate bias = -60V, OctP bias = -18V, and collision and cell helium flow = 4.5 mL/min. Samples were introduced by a peristaltic pump, and taken up at 0.5 rps for 30 s, followed by 30 s at 0.1 rps for signal stabilization. Samples were analyzed by collecting three points across each peak and performing three replicates of 100 sweeps for each element. Data was analyzed using the Agilent Mass Hunter Workstation (version A.01.02). Total iron was quantified by measuring ^56^Fe, which constitutes 91.75% of naturally occurring stable iron isotopes.

**Supplementary Table 1:** Reagents and antibodies

| **Antibody for flow cytometry** | | | | | | | | |
| --- | --- | --- | --- | --- | --- | --- | --- | --- |
| Fluorophore/Antibody | Catalog number | | | | Company | Clone | | Dilution |
| CD16/CD32 Fc Block | 553142 | | | | BD Biosciences | 2.4G2 | | 1:200 |
| BV510 anti-CD45 | 103137 | | | | BioLegend | 30-F11 | | 1:200 |
| PerCP/Cyanine5.5 anti-CD11b | 101227 | | | | BioLegend | M1/70 | | 1:200 |
| APC/Cyanine7 F4/80 | 123117 | | | | BioLegend | BM8 | | 1:200 |
| DAPI | 62248 | | | | ThermoFisher |  | | 1:4000 |
| **Immunofluorescence** | | | | | | | | |
| Hemoglobin | | N/A | | Ciara Shaver Lab | | N/A | | 25 mg/dL |
| DyLight™ 488 NHS Ester | | 46402 | | ThermoFisher | | N/A | | N/A |
| DAPI | | 62248 | | ThermoFisher | | N/A | | 300 μM |
| **Cell Culture Reagents** | | | | | | | | |
| DMEM, high glucose | | | 11965092 | Gibco | | | N/A | N/A |
| Heat Inactivated FBS | | | 10082147 | Gibco | | | N/A | N/A |
| Penicillin-Streptomycin | | | 15140122 | Gibco | | | N/A | N/A |
| Hepes | | | 15630080 | Gibco | | | N/A | N/A |
| L929 conditioned media | | | N/A | This lab | | | N/A | N/A |
| Lipopolysaccharide (LPS) | | | 497693 | ThermoFisher | | | N/A | 1:10000 |
| Interferon-gamma (IFNγ) | | | 485-MI | R&D systems | | | N/A | 1:1000 |
| IL-4 | | | 404-MI | R&D systems | | | N/A | 1:10000 |
| IL-13 | | | 413-MI | R&D systems | | | N/A | 1:5000 |

**Supplementary Table 2:** qPCR reagents and primers

| Reagent/gene | Company | Catalog number | Genomic Map ID |
| --- | --- | --- | --- |
| iScript reverse transcription supermix | BioRad | 1708840 | N/A |
| iQ supermix | BioRad | 1708860 | N/A |
| *Cd163* | ThermoFisher | 4331182 | Mm00474091_m1 |
| *Tfr1* | ThermoFisher | 4331182 | Mm00441941_m1 |
| *Slc11a2* | ThermoFisher | 4331182 | Mm00435363_m1 |
| *Cisd1* | ThermoFisher | 4331182 | Mm00728581_s1 |
| *Hmox1* | ThermoFisher | 4331182 | Mm00516005_m1 |
| *Ftl1* | ThermoFisher | 4331182 | Mm03030144_g1 |
| *Fth1* | ThermoFisher | 4331182 | Mm00850707_g1 |
| *Slc40a1* | ThermoFisher | 4331182 | Mm01254822_m1 |
| *Ncoa4* | ThermoFisher | 4331182 | Mm00451095_m1 |
| *Actb* | ThermoFisher | 4331182 | Mm00607939_s1 |
| *Gapdh* | ThermoFisher | 4331182 | Mm99999915_g1 |

**Supplementary Figure 1:** Body weight and glucose tolerance in female mice. A) Body weight in female CD163^-/-^ and WT littermates on LFD and HFD. B-E) ipGTTs from LFD and HFD female mice after 4 weeks diet (B), 8 weeks diet (C), 12 weeks diet (D), and 16 weeks diet (E). Two-way ANOVA with main effects for time and genotype was used. Data are expressed as mean ± SEM. HFD, high-fat diet; ipGTT, intraperitoneal glucose tolerance test; LFD, low-fat diet.

**Supplementary Figure 2:** Glucose tolerance in male mice after 4, 8, and 12 weeks of LFD or HFD feeding. A-F) ipGTT and ipGTT AUC from CD163^-/-^ and WT littermates on LFD and HFD after 4 weeks (A-B), 8 weeks (C-D), and 12 weeks (E-F). A 2-way ANOVA with main effects for time and genotype was used for panels A, C, and E. A 2-way ANOVA with main effects for diet and genotype was used for panels B, D, and F. # main effect for diet (p<0.05). Data are expressed as mean ± SEM. AUC, area under curve; HFD, high-fat diet; ipGTT, intraperitoneal glucose tolerance test; LFD, low-fat diet.

**Supplementary Figure 3:** Study schematic for hyperinsulinemic clamp. Detailed description of this procedure is found in **Supplementary Methods** section and by Ayala et al., (2).

**Supplementary Figure 4: Peripheral insulin resistance is not impacted by CD163 deficiency in female mice.** A) Body weight was similar between female CD163^-/-^ mice and WT littermates following one-week catheter indwelling in both HFD and LFD fed female mice. B) Insulin concentration increased from basal (t=-10 min) and clamped (t=80-120 min) conditions in all sampled mice, and was greater in the female during the clamp. C) Arterial glucose was measured continuously with target concentration of 150 mg/dL. D) Exogenous GIR was controlled to maintain euglycemia. E) Glucose Rd measured by the sum of exogenous GIR and endogenous glucose Ra. F) Glucose Ra measured during the hyperinsulinemic clamp. G-I) Plasma NEFA measured during the clamp (G), during basal and clamp conditions (H), and NEFA percent suppression from basal to clamp conditions (I). J) A bolus of [^14^C]2DG was infused at 120 min. Mice were then anaesthetized and tissues snap frozen at 155 min to measure ^14^C radioactivity (normalized per 100 g tissue). Unpaired Student’s *t*-test was used to compared differences in CD163^-/-^ and WT littermates in LFD and HFD fed conditions, in panels A, I, and J. A 2-way repeated measures ANOVA was used to assess differences between genotypes over time during the clamp, and conducted in figure panels B-H. Pairwise comparisons were adjusted with Tukey correction. Data are expressed mean ± SEM of 4-7 mice per group. 2DG, 2-Deoxy-D-glucose; Gastroc, gastrocnemius; HFD, high-fat diet; LFD, low-fat diet; NEFA, non-esterified fatty acid; pg AT, perigonadal adipose tissue; Ra, endogenous rate of glucose appearance; Rd, glucose rate of disappearance; SubQ AT, subcutaneous adipose tissue; Vastus L, vastus lateralis. *p<0.05 WT vs. Cd163^-/-^. ^#^p<0.05 main-effect for time during the hyperinsulinemic clamp, basal vs. clamp.

**Supplementary Figure 5:** Loss of CD163 does not affect transcriptome in M0 BMDMs. A) Principle component analysis for M0 BMDMs. B) Volcano plot displaying CD163^-/-^ vs. WT BMDMs. BMDM, bone marrow derived macrophages; PC, principle component.

**Supplementary Figure 6:** Flow cytometry gating strategy, cell number (12-well plates), and viability for hemoglobin uptake assays. A) A live cell gate was applied by forward (FSC) and side (SSC) scatter exclusion, following by doublet exclusion and dead cell exclusion (DAPI^hi^ cells). Median Fluorescence intensity obtained from FITC channel quantified hemoglobin uptake. B-C) Cell number (B) and viability (C) obtained from cells treated with 25 mg/dL hemoglobin in 12-well plates. A 2-way ANOVA was used to assess main effects for genotype (CD163^-/-^ vs. WT) and time (2, 6, 24 h). Data are expressed mean ± SEM of 4 biological samples per group.

**References**

1.	Steele R, Wall JS, De Bodo RC, Altszuler N. Measurement of size and turnover rate of body glucose pool by the isotope dilution method. Am J Physiol. 1956;187(1):15-24. Epub 1956/10/01. doi: 10.1152/ajplegacy.1956.187.1.15. PubMed PMID: 13362583.

2.	Ayala JE, Bracy DP, McGuinness OP, Wasserman DH. Considerations in the Design of Hyperinsulinemic-Euglycemic Clamps in the Conscious Mouse. Diabetes. 2006;55(2):390-7. doi: 10.2337/diabetes.55.02.06.db05-0686.
