## Supplementary figures and images for "Deficiency of the hemoglobin-haptoglobin receptor, CD163, worsens insulin sensitivity in obese male mice"

### Supplementary Figure 1.tif

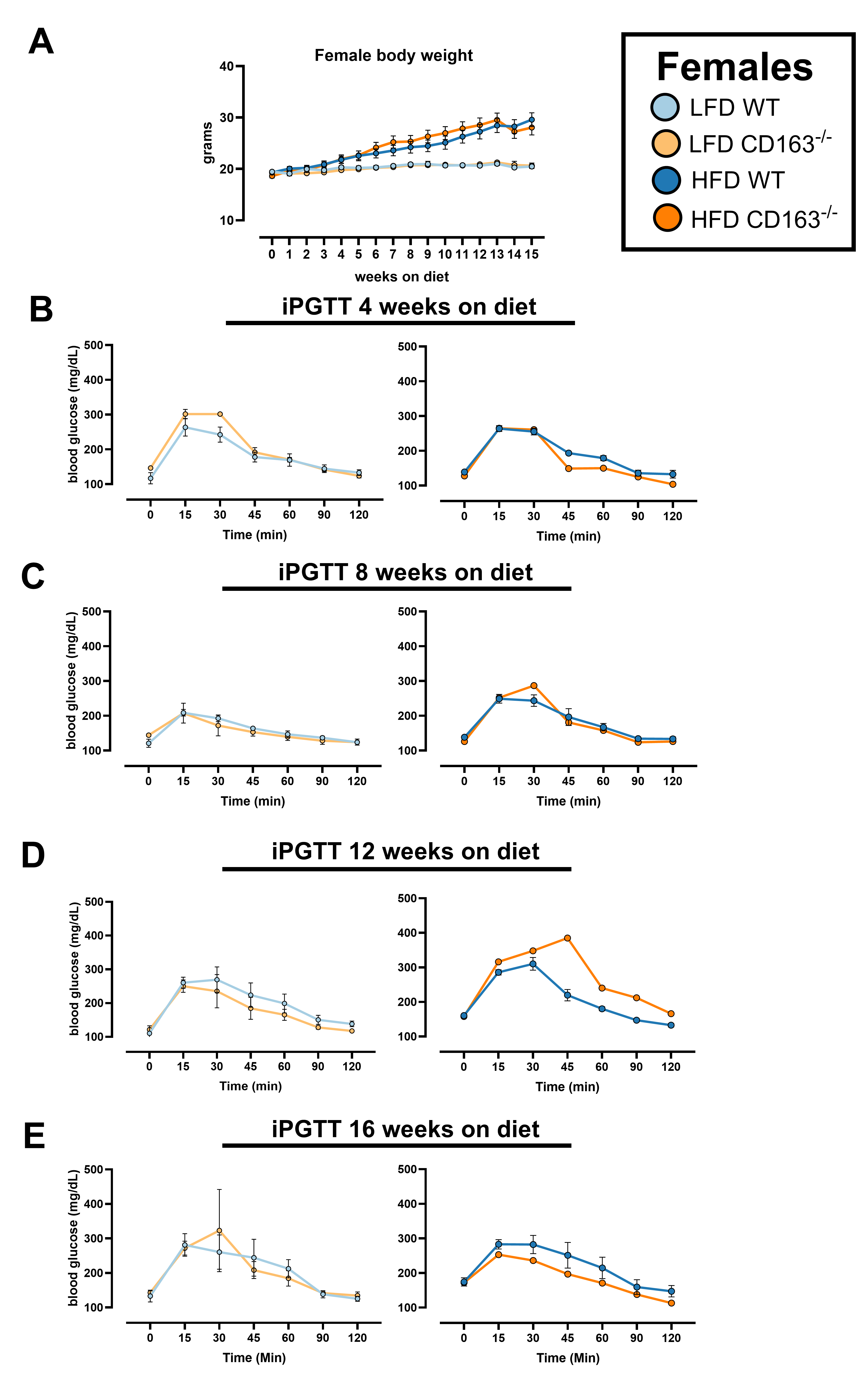

### Supplementary Figure 2.tif

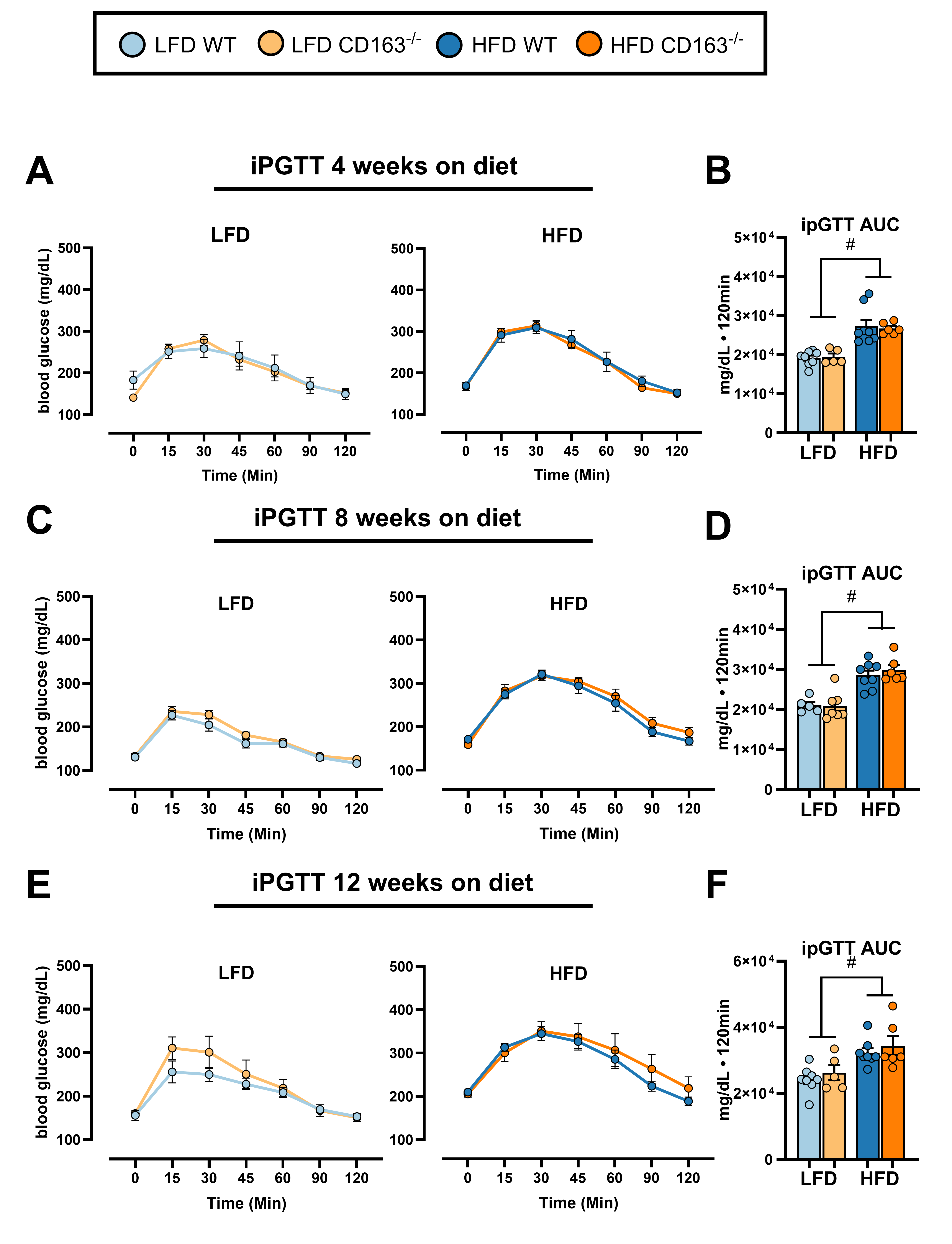

### Supplementary Figure 3.tif

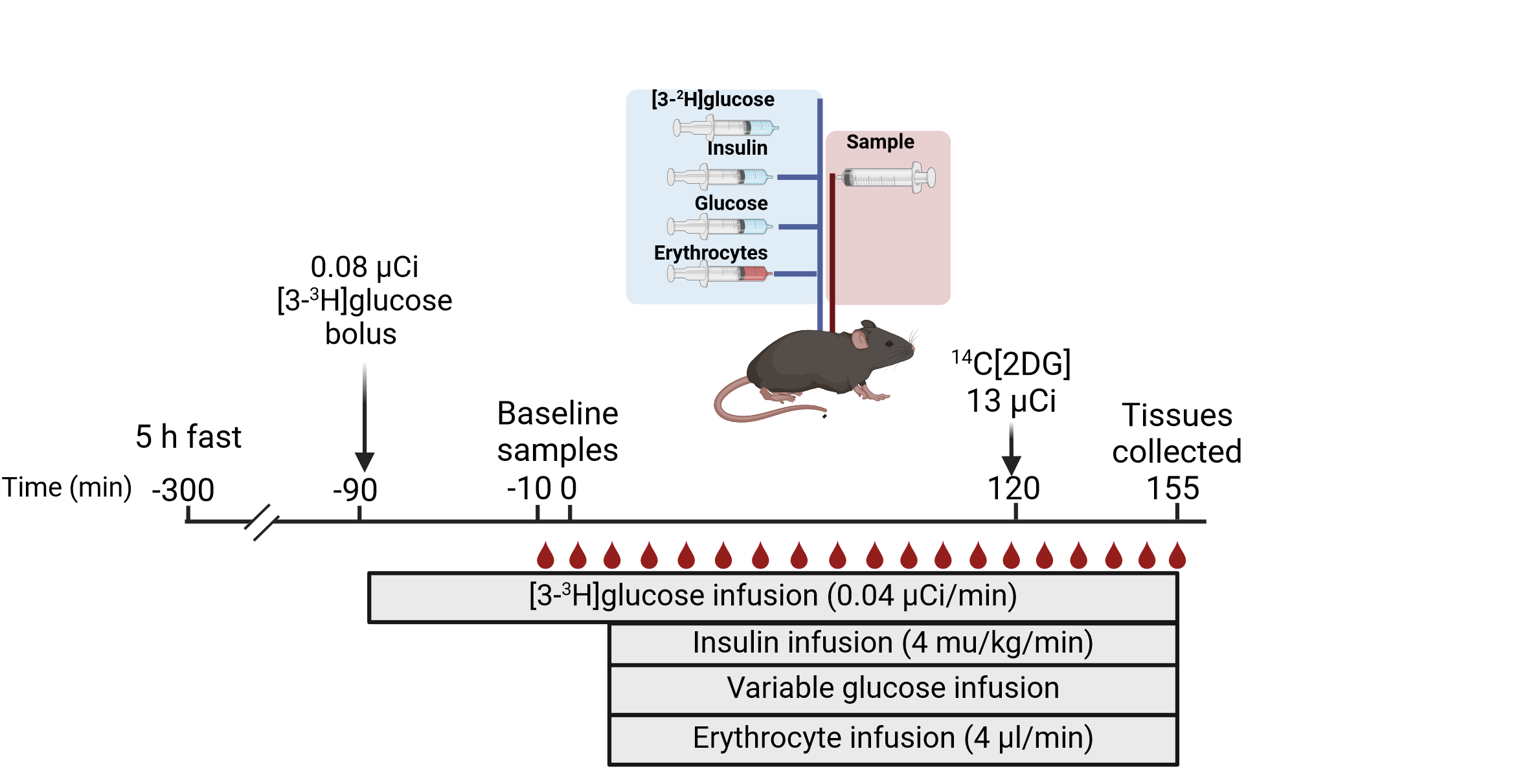

### Supplementary Figure 4.tif

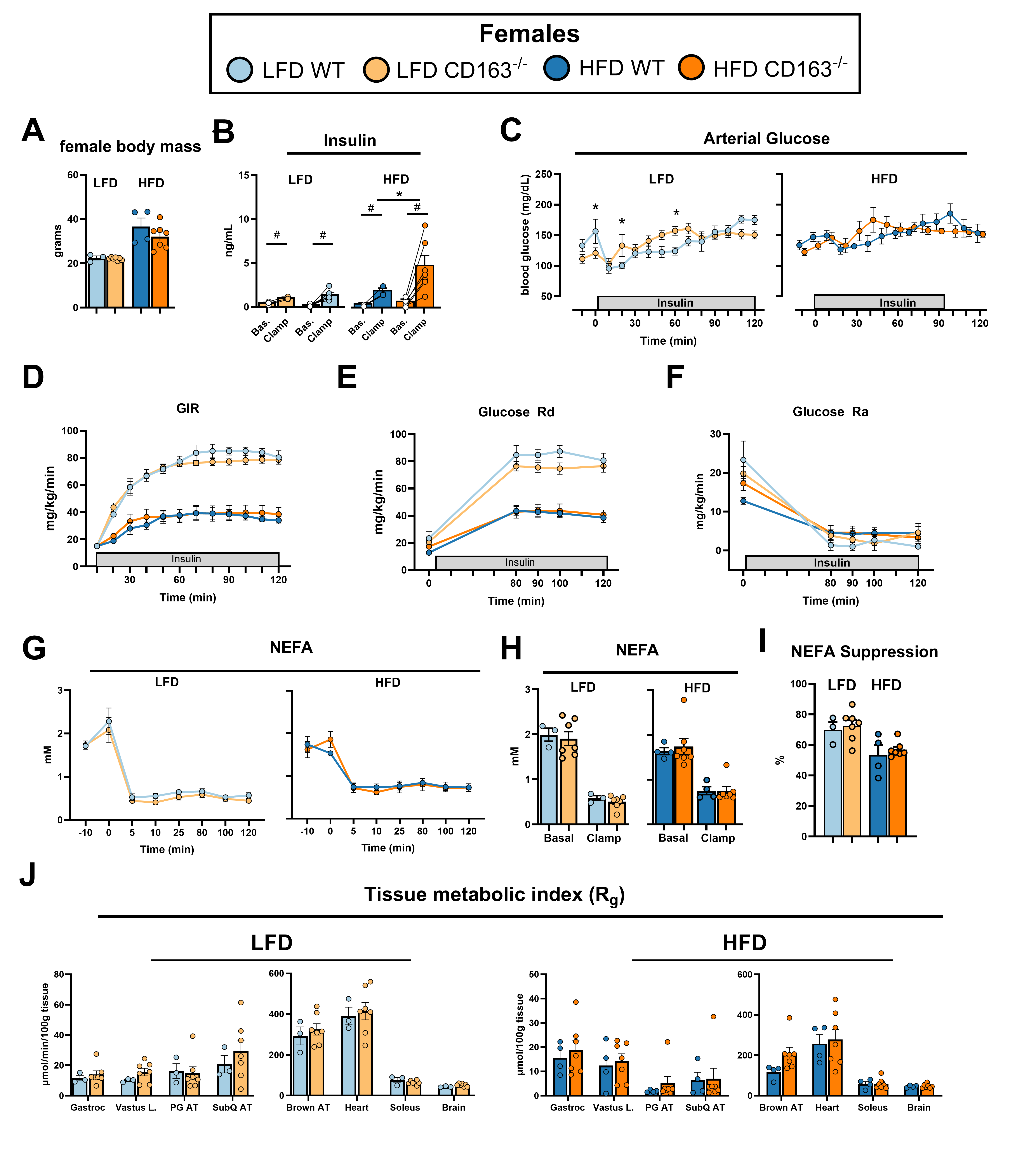

### Supplementary Figure 5.tiff

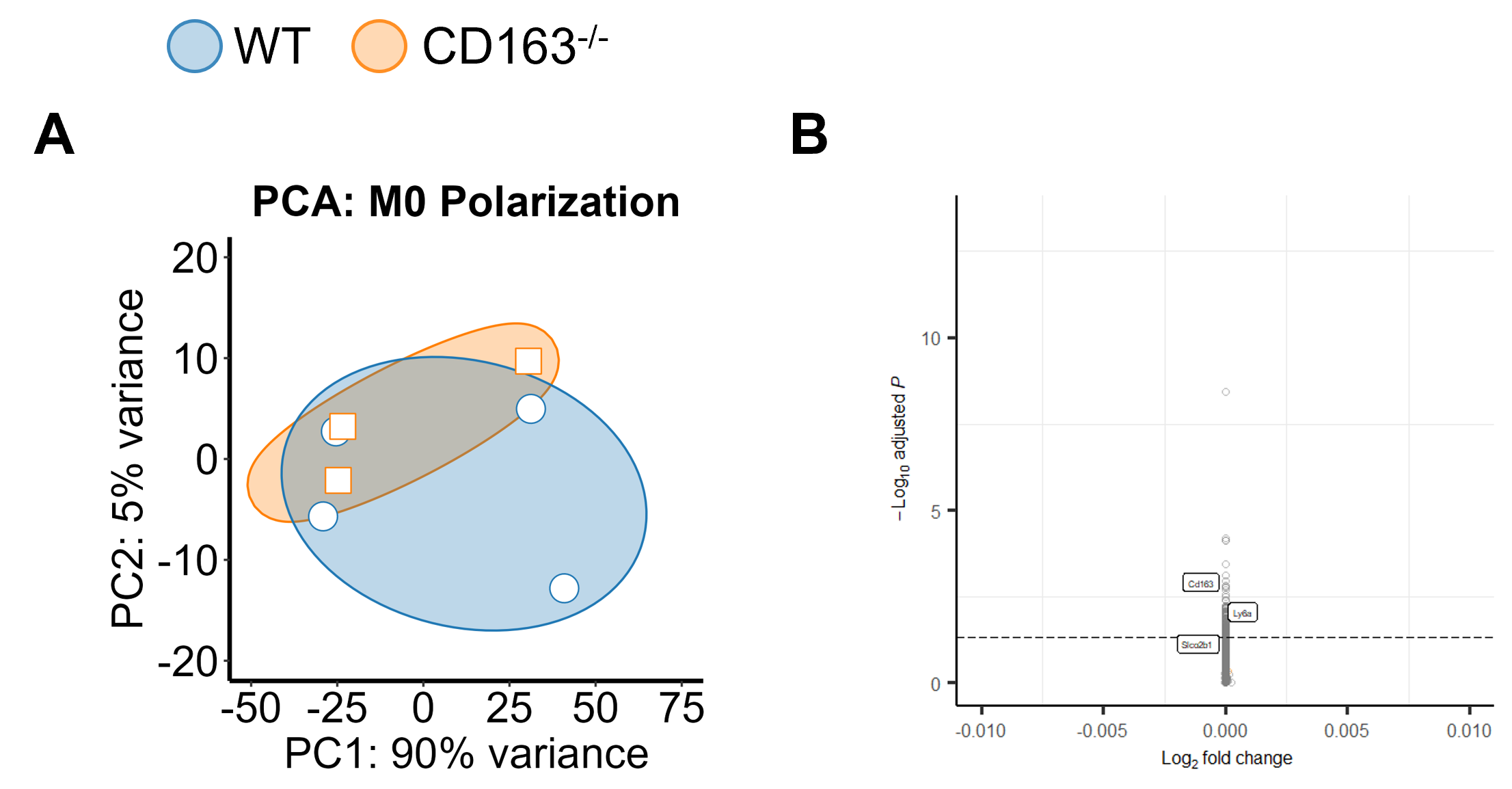

### Supplementary Figure 6.tif

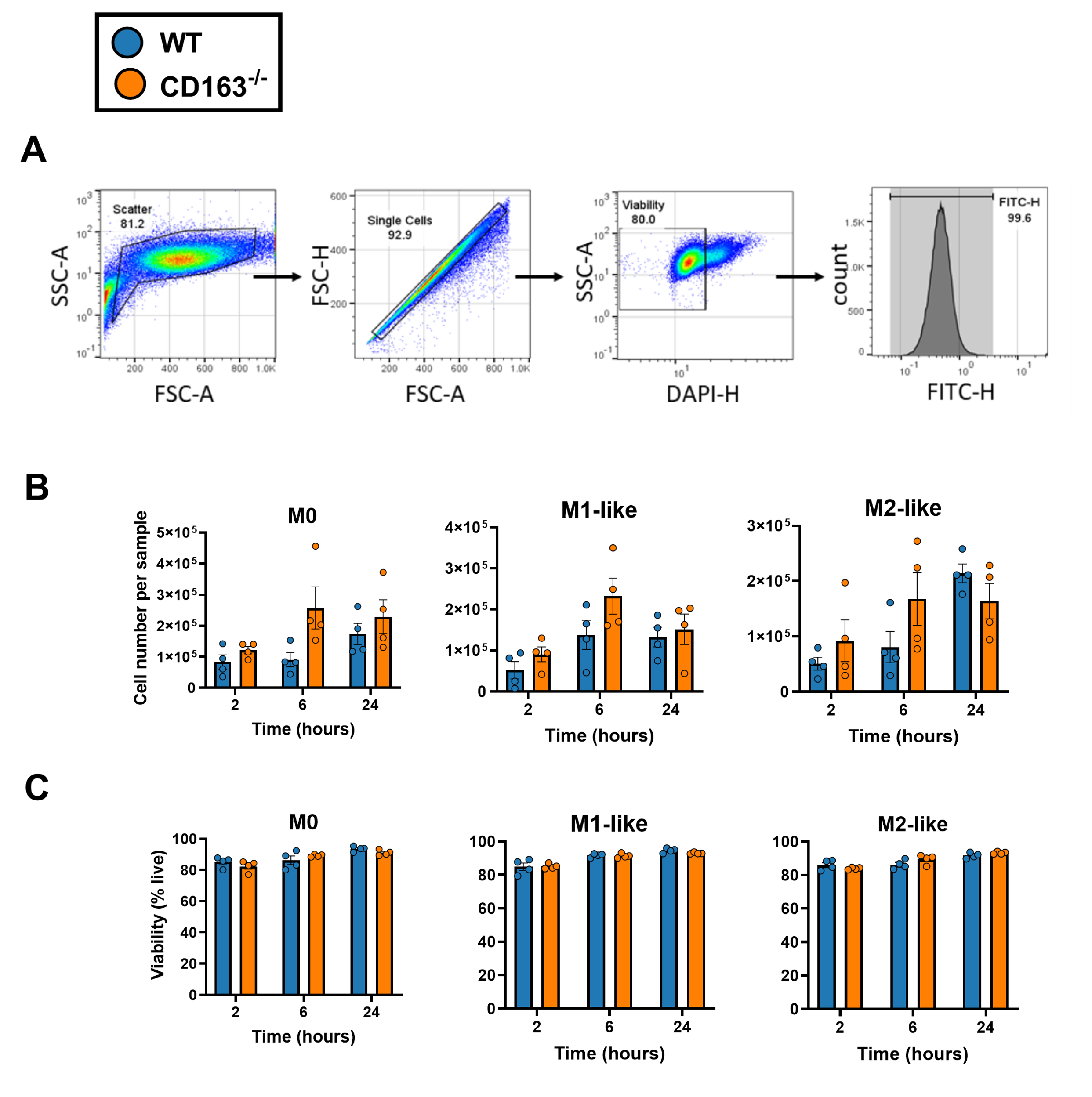
